# The genetic repertoire of the deep sea: from sequence to structure and function

**DOI:** 10.64898/2026.02.03.703410

**Authors:** Yang Guo, Zongan Wang, Denghui Li, Lele Wang, Hongxia Lan, Fei Guo, Ziyu Zhao, Zhenjun Liu, Liang Meng, Xuechun Shen, Minxiao Wang, Weishu Zhao, Weijia Zhang, Chaodi Kong, Liuxin Shi, Ying Sun, Inge Seim, Aijun Jiang, Kailong Ma, Zidong Su, Nannan Zhang, Qianyue Ji, Junyi Chen, Ke Chen, Chen Qi, Baitao Li, Beibei He, Yuqian Liu, Jiayong Zhou, Yue Zheng, Huan Zhang, Yinzhao Wang, Mo Han, Tao Yang, Jiawei Tong, Yulu Zhang, Zhijie Wang, Xiaokai Xu, Jiayu Chen, Yue Liu, Haixin Chen, Tao Zeng, Xiaofeng Wei, Chaolun Li, Huanming Yang, Bo Wang, Xin Liu, Changwei Shao, Wenwei Zhang, Ying Gu, Xiang Xiao, Xun Xu, Jian Wang, Thomas Mock, Guangyi Fan, Yuxiang Li, Shanshan Liu, Yuliang Dong

**Author notes:** Correspondence (Y. D.), (S. L.), (Y. L.), (G. F.), (T. M.). These authors contributed equally to this work (Yang Guo, Zongan Wang, Denghui Li, Lele Wang, Hongxia Lan).

## Abstract

The deep sea as the largest and maybe most hostile environment on Earth is still underexplored especially regarding its genetic repertoire. Yet, previous work has revealed significant habitat-specific deep-sea biodiversity. Here, we present an integrated deep-sea genetic dataset comprising 502 million nonredundant genes from 2,138 samples and 2.4 million predicted structures, and used it to link specific protein structures with genetic variants associated with life in the deep sea and to assess their biotechnology potential. Combining global sequence analysis with biophysical and biochemical measurements revealed unprecedented sequence diversity, yet substantial structural conservation of proteins. Especially proteins involved in replication, recombination, and repair were identified to be under rapid evolution and with specialized properties. Among these, a structurally divergent helicase exhibited advantages in controlling nanopore sequencing speed. Thus, our work positions the deep sea as a unique evolutionary engine that generates and hosts genetic diversity and bridges genetic knowledge with biotechnology.

## Introduction

The deep sea, particularly the regions below the mesopelagic zone at depths exceeding 1,000 meters, represents Earth’s largest and at the same time least explored biome^1, 2^. It hosts habitats such as hadal trenches, hydrothermal vents, and methane seeps occupied with diverse life forms adapted to survive these harsh environmental conditions from near freezing temperatures (< 4°C)^2, 3^ to high temperatures above 100°C at hydrothermal vents^4^. These temperature extremes are accompanied by hydrostatic pressure of up to ca. 110 MPa in the Mariana Trench^3^, permanent darkness and limited oxygen availability^2^. Pioneering explorations in the 1950s^5^ and 1960s^6^, followed by the discovery of deep-sea chemosynthetic ecosystems in the 1970s^7^, advanced the study of deep-sea life^8, 9^. Studies have reported significant biodiversity, especially for microbiomes at hydrothermal vents^9, 10^ and methane seeps^11, 12^. Furthermore, recent initiatives like the Mariana Trench Environment and Ecology Research (MEER) project report significant diversity of hadal-trench microbiomes including their metabolic pathways^13, 14^. Despite all these research efforts, our knowledge about the genetic repertoire of the deep sea is still elusive. Yet, even this limited understanding may be on par with our knowledge of more accessible, smaller ecosystems on Earth.

The development of genetic resources based on large-scale metagenomic datasets can advance biotechnological and biomedical applications^15^. The deep sea has long been considered a valuable source of genetic resources with significant potential for diverse applications to meet the demands of human societies^16, 17^. The extreme environment of the deep-sea such as elevated hydrostatic pressure, toxic metals, and radicals, can damage nucleic acid and therefore might affect core biological processes including replication and transcription of nucleic acids^18, 19^. Thus, proteins involved in replication, recombination, and DNA repair from deep-sea microorganisms may have evolved unique protein structures and therefore enzymatic characteristics such as shown by the DNA polymerases isolated from *Thermococcus litoralis* and *Thermococcus thioreducens*^20, 21^. Consequently, these deep-sea proteins are potentially invaluable for molecular diagnostics, molecular biological research, and DNA sequencing.

Recent advancements in deep-sea sampling and AI-based protein structure prediction tools have laid the basis for comprehensively characterizing the deep-sea genetic diversity^22–24^. To address this overarching aim, we integrated 2,138 metagenomic datasets (1,194 from the MEER project^14^ and 944 from public databases) to analyze the sequence diversity and the habitat-dependent distributions of deep-sea unique gene families, and revealed that genes involved in replication, recombination, and repair are rapidly evolving in the deep-sea. In addition, we predicted 2.4 million deep-sea protein structures, most of which exhibit homologies with the AlphaFold Protein Structure Database (AFDB)^25, 26^. However, we also identified 392 novel protein domains that appear to be exclusive to deep-sea microbes. Furthermore, we illustrated how structure-homology-based methods can aid the discovery of novel deep-sea enzymes (e.g., helicase) with biotechnological potential. Therefore, our study deepens our knowledge of the ecological and evolutionary divers of the deep-sea genetic diversity and provides a framework for harnessing this largely untapped biological resource.

## Results

### The genetic diversity of deep-sea microbiomes

To characterize the genetic diversity of deep-sea microbiomes (here at depths ≥ 1,000 m), we compiled the metagenomic data of 2,138 samples (1,194 from the MEER project^14^ and 944 deposited at NCBI) spanning global deep-sea environments (Figures 1A and 1B), including water (364), sediment (1,617), and others such as biofilms and rocks (157) (Figure S1A and Table S1). We categorized these samples into four groups according to their habitat type: hydrothermal vents (305), methane seeps (167), hadal ecosystems (1,260) and other deep-sea ecosystems (406) (Figure 1A and Table S1). From these datasets, we predicted 1.9 billion protein-coding genes and clustered them at > 95% sequence identity, generating a nonredundant Deep-Sea Gene Catalog (DSGC) of 502 million unigenes (Figures 1C and S1B). While 58.44% of the unigenes had incomplete open reading frames, 86.27% of these incomplete genes were merged into gene clusters containing at least one complete sequence (> 20% amino-acid identity) (Figure S1C). DSGC captured 69.19% of high-quality deep-sea metagenomic reads (Figure S1D), which rate was consistent with the global ocean microbial reference gene catalog (OM-RGC, v2)^27^, indicating the effectiveness in covering the majority of gene-encoding metagenomic data. Interrogating public databases revealed the presence of significant sequence novelty in the DSGC: 51.13% and 73.37% of genes lack functional annotations using EggNOG^28^ and KEGG^29^ respectively, and 53.79% were taxonomically uncategorized at the domain level (Figure 1D). Against two additional and global gene catalogs: OM-RGC (representing the global upper ocean)^27^ and TSGC (the global topsoil gene catalog, representing global topsoil)^30^, DSGC exhibited the highest diversity (as indicated by number of effective sequences, N ^31^) in 80.43% of all detected Pfams (Methods, Figure 1E). Furthermore, clustering DSGC sequences with OM-RGC and TSGC at > 20% amino-acid identity revealed that deep-sea genes formed over 58 million unique clusters (Figure 1F), and 18 million deep-sea unique clusters remained when excluded singletons from all clusters (Figure S1E). This pattern remained robust in controlled analyses accounting for both dataset scale and taxonomic composition (Methods, Table S2, S3). When clustered with sequences of the Global Ocean Gene Catalog^32^ at > 20% amino-acid identity, DSGC contributed 157.13 million additional sequences as part of 55 million unique gene clusters, expanding the genetic diversity of the Global Ocean Gene Catalog^32^ (308.57 million non-redundant sequences) by 50.92% (Figure 1G). These results underscore the sequence diversity and uniqueness of deep-sea microbial genes, and raise the question as to how they have evolved.

**Figure 1.**
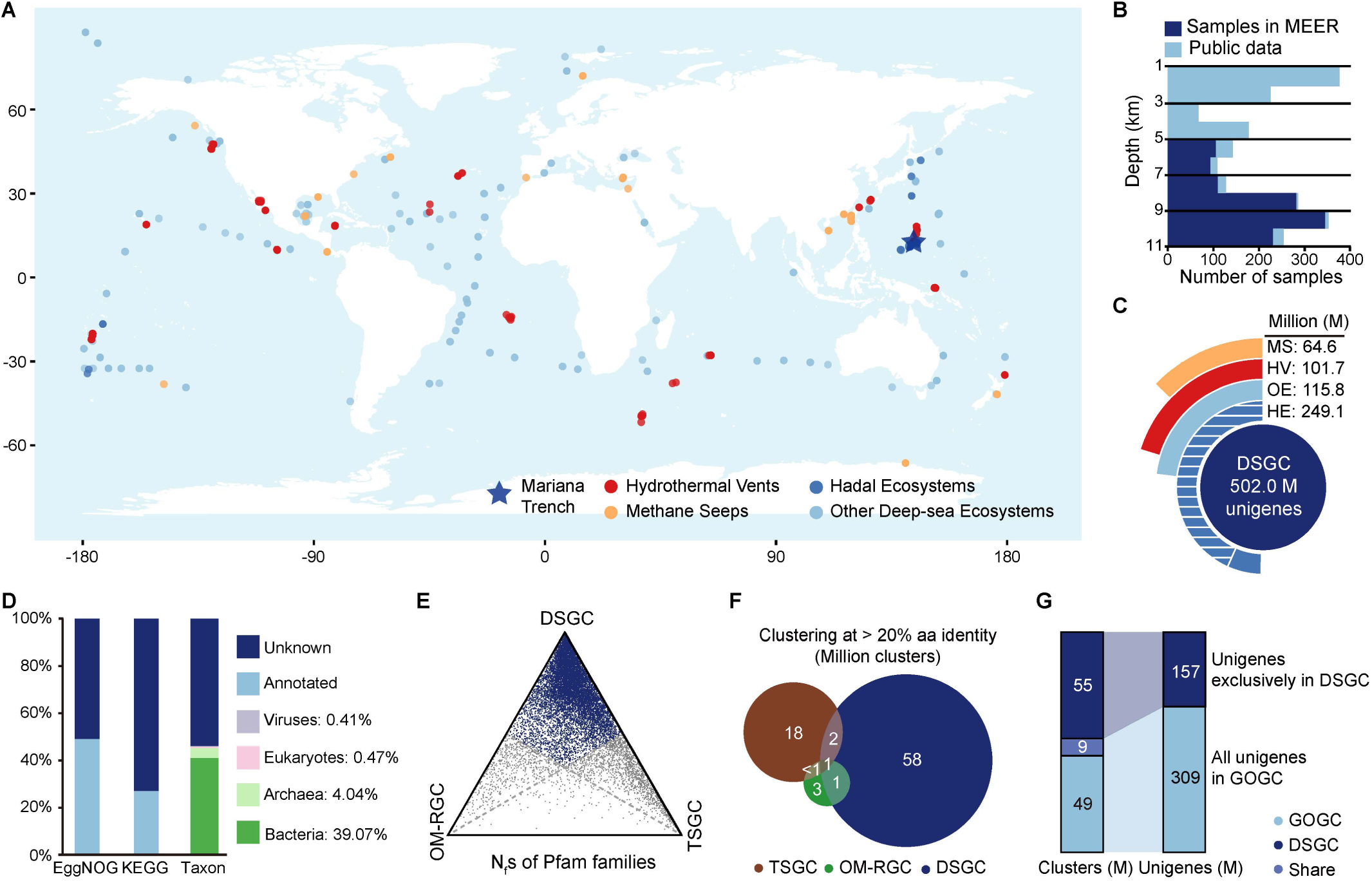
Integrated deep-sea microbial gene catalog (DSGC). **(A)** Geographic distribution of the 2,138 deep-sea samples in DSGC. Samples were classified into four habitat types: hydrothermal vents, methane seeps, hadal ecosystems, and other deep-sea ecosystems. **(B)** Depth breakdown of samples used in this study. Samples contributed by the MEER project are highlighted in dark blue. **(C)** Total gene count and breakdown of gene number by habitat in DSGC. The shadowed zone in hadal ecosystem indicates the unigenes contributed by the MEER project. HV: hydrothermal vents, MS: methane seeps, HE: hadal ecosystems, and OE: other deep-sea ecosystems. M: million unigenes. **(D)** Functional and taxonomic annotation of DSGC. **(E)** Ternary plot showing the proportional relations of the number of effective sequences (Nf) for each Pfam family. Each axis represents the proportion of a family’s Nf found in one catalog relative to the total across DSGC, OM-RGC, and TSGC. A single dot represents one Pfam family; its position encodes the triple proportion (summing to 100%). Blue dots indicate families where the DSGC’s Nf is the highest among the three catalogs. **(F)** Venn diagram of unique and shared gene clusters (at > 20% amino-acid identity, in millions) among DSGC, OM-RGC, and TSGC. **(G)** Unique and shared gene clusters (in millions) between DSGC and the Global Ocean Gene Catalog (GOGC), and the number of unigenes (in millions) in exclusively DSGC gene clusters.

### Habitat-dependent distribution of deep-sea microbial genes

The extreme habitats of the deep sea drive microbial adaptations^4, 13^ and thus may cause either the evolution of entirely novel genes or variants of genes with known function. As for the latter, categorizing genes according to their orthologous groups (COGs) revealed that the COG-L category (“replication, recombination, and repair”) was the most diverse category in the annotatable part of DSGC (10.42%) (Figure S2A). This proportion was greater than those in the upper ocean (OM-RGC: 6.24%) and topsoil (TSGC: 7.09%) (Figure S2A), and the high proportion in DSGC was not likely favored by certain phyla with more diverse COG-L genes (Figure S2B), suggesting that the COG-L category genes in deep-sea microbiomes may have undergone a significant level of divergence. Additionally, DSGC covered 93.60% (731) of the 781 COG terms from the L category, while the numbers for OM-RGC and TSGC were 76.57% (598) and 84.38% (659), respectively. Even when restricted to the shared 598 COG-L terms among these three catalogs, DSGC still contained the largest number of non-redundant genes (26.87 million), exceeding those in OM-RGC (1.92 million) and TSGC (8.24 million). Furthermore, deep-sea microbial genomes contained significantly higher proportions of COG-L category genes compared to the microbes from the upper ocean (Figure S2C). Considering the significant divergence of deep-sea COG-L genes, together with their potential role in coping with environmental stressors that can damage nucleic acids^18, 19^, we focused on the COG-L category genes as a test case.

By clustering all COG-L category genes from the three global gene datasets at > 20% amino-acid identity^33^, we grouped these homologous into 728,097 gene clusters, with 243,218 clusters only identified in the deep sea (denoted as “excl-DS-cluster”, when compared with the OM-RGC and TSGC) (Figure 2A). A substantial portion (33.81%) of excl-DS-clusters were indeed rare (detected in < 1% of samples) (Figure 2B). This finding, though focused on a specific COG category, is consistent with the Global Microbial Gene Catalog, which proposed that habitat-specific rare genes can arise through neutral evolution with limited selection^33^. At the same time, 43.40% of excl-DS-clusters (105,558 clusters) were more prevalent in the deep sea (detectable in > 100 samples) (Figure 2B), suggesting a complex distribution pattern potentially linked to microbial survival strategies.

**Figure 2.**
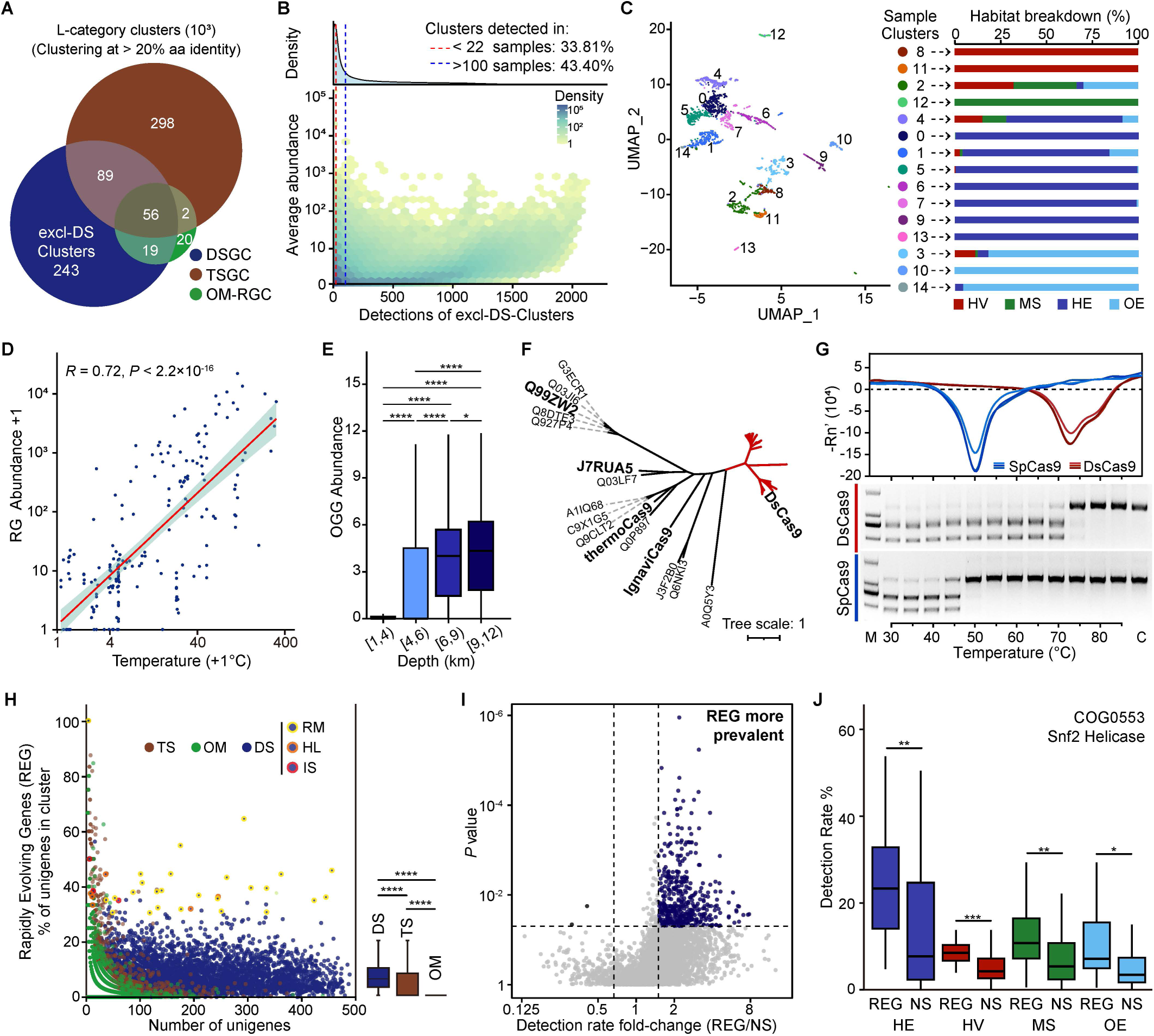
Distribution and evolution of COG-L genes in deep-sea environments. **(A)** Venn diagram of unique and shared COG-L gene clusters (at > 20% amino-acid identity) among DSGC, OM-RGC, and TSGC. excl-DS-clusters: exclusive deep-sea gene clusters. **(B)** Detections and average abundances of excl-DS-clusters. The distribution of excl-DS-Cluster detections is shown by the curve at the top. The dashed red and blue lines mark the thresholds of 22 (1% of all samples) and 100 detections. **(C)** UMAP visualization of the abundances of enriched and prevalent excl-DS-clusters from 2,138 samples and habitat breakdown of samples in each cluster. **(D)** Abundance of a reverse gyrase (*RG*, COG1110) cluster in samples at different temperature. Shading of the fitted line shows 95% confidence interval. **(E)** Abundances of an 8-oxoguanine DNA glycosylase (*OGG*, COG4047) gene cluster in samples at different depths. **(F)** Phylogenetic tree of members in the hydrothermal vent-specific *Cas9* (COG3513) cluster (red branches), two previously reported thermostable *Cas9s* (*thermoCas9* and *IgnaviCas9*), and well-annotated *Cas9s* from Swiss-Prot^115^, including *SpCas9* (Q99ZW2) and *SaCas9* (J7RUA5). **(G)** Melt-curves and *in-vitro* genome editing activities of DsCas9 and SpCas9 proteins under different temperatures. M: marker, C: blank control. **(H)** Frequencies of rapidly evolving genes (REGs) in each gene cluster. Frequencies were calculated separately for genes from DSGC (DS), OM-RGC (OM), and TSGC (TS). The boxplot on the right represents the distributions of the rapidly evolving frequencies without outliers. RM: genes related to restriction-modification, HL: helicases, IS: topoisomerases. **(I)** Volcano plot indicating the fold-change of detection frequencies of REGs and non-rapidly evolving genes (gene with no significant probability, NS) in each gene cluster. Blue dots denote clusters where the detection frequencies of REGs were significantly higher (*P* < 0.05, fold-change > 1.5) than NSs across the 2,138 deep-sea samples. **(J)** Detection rates of REGs and NSs from a *Snf2* helicase cluster in the four habitat groups. **(C, J)** the abbreviations are as follows: HV: hydrothermal vents, MS: methane seeps, HE: hadal ecosystems, and OE: other deep-sea ecosystems. **(E, H, J) s**ignificances of paired comparisons were calculated using a two-tailed Wilcoxon test (\**P* < 0.05, \*\**P* < 0.01, \*\*\**P* < 0.001, \*\*\*\**P* < 0.0001).

To investigate the ecological drivers of these prevalent excl-DS-clusters, we clustered 2,138 deep-sea samples based on their gene abundance profiles. For our primary analysis, we used the relative abundances (Methods) of only those prevalent (detection > 100) excl-DS-clusters that were enriched for COG functional terms. This analysis revealed a robust association between deep-sea habitat types and gene distribution (Cramer’s *V* = 0.74) (Figure 2C), with the significance retained even for all excl-DS-clusters with more than 100 detections (Cramer’s *V* = 0.70) (Figures S2D and S2E).

This habitat-specific distribution is exemplified by several excl-DS-clusters whose functions aligned with the harsh physicochemical regimes in the deep sea. For instance, an excl-DS-cluster annotated as reverse gyrase (COG1110), with members belonging to both bacterial and archaeal phyla (Figure S2F), was significantly enriched at hydrothermal vents (53.43-fold, *P* < 2.22×10^-16^). Given that high temperature is a defining and extreme factor of these marine habitats, we found that their abundance correlated well with ambient temperatures (*R* = 0.72, *P* < 2.22×10^-16^) (Figure 2D). Reverse gyrases can generate positive supercoils of DNA to maintain the integrity of DNA for facilitating protein-DNA interactions (e.g., transcription) under high temperatures^34^. Similarly, *Cren7*, a gene coding chromatin protein that constrains negative DNA supercoils^35^, had significantly higher abundance (2,770.59-fold, *P* < 2.22×10^-16^) at hydrothermal vents. In hadal zones, a bacterial-archaeal 8-oxoguanine DNA glycosylase cluster (*OGG*, COG4047) (Figure S2G), was significantly enriched (7.16-fold, *P* < 2.22×10^-16^) and still was significantly more abundant in the deeper oceans than zones shallower than 4,000 m (Figure 2E). This enzyme class is responsible for antioxidation via repairing oxidized guanine bases in DNA^36^. Antioxidation is an important mechanism for microbes to survive from high hydrostatic pressure^37, 38^, and signals of enhanced capabilities in antioxidation and DNA repair have been observed in hadal lives and high hydrostatic pressure-tolerant mutants^39–41^.

Furthermore, among the deep-sea unique gene clusters, we identified a *Cas9* variant (COG3513) specifically enriched (99.53-fold, *P* < 2.22×10^-16^) in hydrothermal vents ecosystems (*DsCas9*). Phylogenetic analysis revealed a substantial sequence divergence between *DsCas9*, thermostable *Cas9s* (*thermoCas9*^42^ and *IgnaviCas9*^43^), and other well-characterized homologs (*SaCas9*^44^ and *SpCas9*^45^) (Figure 2F). To probe the thermal adaptation of *DsCas9*, we expressed a hydrothermal-vent-derived DsCas9 protein (sampled at 318°C^46^) and observed a melting temperature of 73°C, which is significantly higher than that of the commonly used SpCas9 (49°C) (Figure 2G). We predicted its guide RNA sequence and verified its human genome editing activity (Figures S2H and S2I). This DsCas9 was active at temperatures up to 75°C, surpassing SpCas9 (50°C), whereas DsCas9 showed the best activity in approximately 100 mM NaCl (Figures 2G and S2J). Thus, our functional characterization confirms that the DsCas9 protein, sourced from deep-sea hydrothermal vents, possesses high thermostability congruent with the extreme thermal regimes of its native habitat.

Collectively, these examples demonstrate how distinct deep-sea environments can harbor genes with adaptive traits.

### Evolutionary forces and their impact on the genetic diversity of the deep sea

Genes from the deep sea that had at least 20% amino-acid sequence identity to homologs in other sequence databanks (Figure 2A) were used for evolutionary analyses, and these genes also exhibited significant sequence diversities when compared to their homologs from the upper ocean and topsoil (Figure S2K).

We first investigated the selective pressure in the deep-sea environments on these genes with two independent approaches, including per-branch selection tests within each cluster and estimation of nucleotide diversity and pN/pS ratios. The selection tests revealed that deep-sea microbes harbor a significantly higher proportion (median: 6.06%, IQR: 3.12–10.26%) of rapidly evolving genes (REGs) compared to topsoil (median: 0%, IQR: 0–8.33%) and upper-ocean microbes (the upper quartile: 0%) (Methods, Figures 2H and S2L). The latter analysis confirmed the existence of elevated non-synonymous substitution levels (*P* < 2.2×10^-16^) with maintained nucleotide diversity (*P* = 0.073) in those REGs (Figures S2M and S2N). Specifically, gene clusters related to restriction-modification, topoisomerases, and helicases were among the clusters with most frequently (> 30%) observed rapid evolution (Figure 2H), implying the crucial role of these gene families in the adaptation of microbes to the deep sea. For instance, consistent with the identification of a hydrothermal vent-specific monomeric topoisomerase, the reverse gyrase (Figure 2D), we found three topoisomerases (belongs to topoisomerase IB and IIA) among the top 50 rapidly evolving gene clusters (Figure 2H), suggesting the importance of maintaining DNA integrity under high pressure and/or elevated temperatures.

The combination of higher frequencies of rapid evolution and elevated pN/pS (while mostly under purifying selection at the gene level, i.e., gene-wide pN/pS < 1) suggests that these genes are evolving under distinct selective regimes. Our observation of maintained nucleotide diversity argues against a pervasive relaxation of purifying selection, which would be expected to increase diversity^47–49^. This pattern, instead, supports localized positive selection acting on specific sites. Such a mechanism can drive rapid evolution and elevate pN/pS while remaining compatible with maintenance of gene-wide nucleotide diversity^50, 51^. This maintenance of diversity, rather than a significant decrease typically observed as a consequence of a selective sweep, is plausible, if, the selective sweeps either took place in the distant evolutionary past or the signals are diluted by the surrounding genetic loci which evolved under strong purifying selection^52, 53^.

We partially addressed this issue by comparing REGs to non-rapidly evolving genes (with no significant probabilities of rapid evolution) within the same clusters. The REGs exhibited higher detection frequencies across deep-sea samples in 81.66% of the analyzed clusters (3,517/4,307), with 14.16% (498/3,517) showing significant higher detection frequencies (Figure 2I). For instance, REGs in a *Snf2* helicase (COG0553) cluster were significantly more prevalent across different deep-sea habitats (Figure 2J).

Aligning with the pattern of maintained nucleotide diversity (Figure S2N), our ecological prevalence data therefore offers a complementary perspective to assess these evolutionary drivers. The pervasive deletion bias in prokaryotic genomes creates a constant pressure for the loss of non-essential DNA^54–56^. This makes the widespread prevalence of REGs difficult to reconcile with a model of relaxed selection alone, as such genes would be prone to stochastic loss. While hitchhiking can cause the occasional retention of such genes, the adaptive benefit conferred by positive selection provides a more direct and parsimonious explanation for their systematic success. Therefore, the observed ecological success indicates that while relaxed selection may influence sequence-level variability, positive selection of specific genetic loci for adaptive evolution likely plays a role in the maintenance and successful expansion of these genetic variants across diverse marine habitats.

### Structural homology of deep-sea proteins

Due to the significant divergence of genes in the deep sea, sequence-based annotation and bioprospecting are challenging. To overcome this challenge, we constructed a deep-sea protein fold catalog (DSFC), because previous research has demonstrated that the protein structure exhibits greater evolutionary conservation compared to their underlying gene sequences^57–59^.

We focused on 2.4 million gene clusters (≥ 30 members per cluster at > 20% amino-acid identity) and used ESMFold^23^ to predict the structures of the representative proteins (Methods) of these clusters. This resulted in 2.4 million deep-sea protein structures, which included 1.1 million high-confidence predictions and represented nearly 70% (340.2 million) of the proteins in DSGC (Figures 3A, S3A and S3B). We showed that the deep-sea protein structures predicted by ESMFold^23^ were highly convergent with those predicted by AlphaFold2^22^, suggesting consistency between both methods (Figure S3C). For subsequent analyses, we only used high-confidence structures.

**Figure 3.**
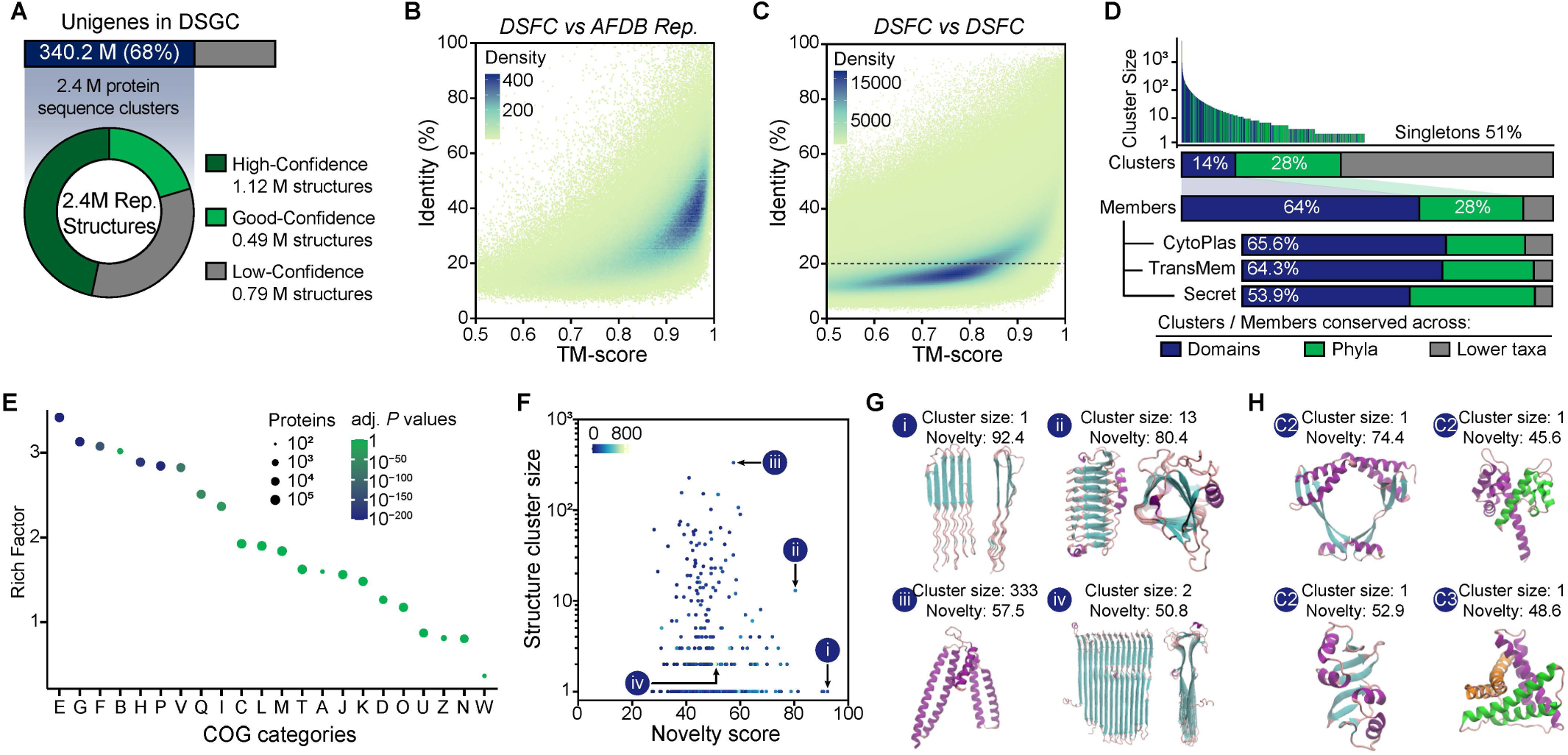
Structural homologies of deep-sea proteins and novel domains in DSFC. **(A)** The deep-sea protein fold catalog (DSFC). The catalog contains approximately 2.4 million structures, including 1.12 million high-confidence (mean pLDDT > 0.7 and pTM > 0.7) structures, 0.82 million with good-confidence (mean pLDDT > 0.5 and pTM > 0.5), and 0.46 million with low confidence (mean pLDDT <= 0.5 or pTM <= 0.5). **(B)** Distribution of structural and sequence similarity between proteins from DSFC and the clustered AFDB (AFDB Rep.). **(C)** Distribution of structural and sequence similarity between proteins from DSFC. **(D)** Structural conservation between deep-sea archaea and bacteria. Taxonomically categorized structures in DSFC were grouped into cytoplasmic (CytoPlas), trans-membrane (TransMem) and secreted (Secret) proteins. **(E)** Enrichment of COG categories with structures conserved across deep-sea archaea and bacteria. Significances were calculated using the Fisher’s exact test. The rich factor was the ratio between structure numbers of conserved inter- (conserved across domains) and intra-domains (conserved across lower taxa within either Archaea or Bacteria). Abbreviations are sorted according to the horizontal axis: E, Amino acid transport and metabolism; G, Carbohydrate transport and metabolism; F, Nucleotide transport and metabolism; B, Chromatin structure and dynamics; H, Coenzyme transport and metabolism; P, Inorganic ion transport and metabolism; V, Defense mechanisms; Q, Secondary metabolites biosynthesis, transport and catabolism; I, Lipid transport and metabolism; C, Energy production and conversion; L, Replication, recombination and repair; M, Cell wall/membrane/envelope biogenesis; T, Signal transduction mechanisms; A, RNA processing and modification; J, Translation; ribosomal structure and biogenesis; D, Cell cycle control, cell division, chromosome partitioning; K, Transcription; O, Posttranslational modification, protein turnover, chaperones; U, Intracellular trafficking, secretion, and vesicular transport; Z, Cytoskeleton; N, Cell motility; W, Extracellular structures. **(F)** Novelty of novel domains identified in DSFC (*n* = 392). Dots are colored by the size of structural cluster to which the protein containing the novel domain belong. Novelty scores were predicted following the TED protocol^65^. (i) to (iv) correspond to labels shown in **G.** **(G)** Examples of novel domains. The novel domains with the highest novelty score (i), in the largest structural cluster (iii), and with the most residues (iv) were selected for visualization. **(H)** Examples of novel domains with internal symmetry. C2 and C3 stand for twofold and threefold axial symmetry, respectively.

We found that 97.97% of high-confidence structures had homologous structures (TM-score > 0.5^60^) in AFDB^26, 61^ and 80.25% with PDB^62^ entries (Figures 3B and S3D). Pairwise comparisons further revealed widespread structural conservation (TM-score > 0.5) among deep-sea proteins, despite substantial sequence differences (< 20% amino-acid identity) (Figure 3C). Furthermore, clustering 759,874 protein structures with associations to known taxa identified 64% cross-domain structures between bacteria and archaea (Figure 3D), including proteins involved in transport and metabolism of amino acids, nucleic acids, cofactors, and metal ions (Figure 3E). Interestingly, secreted proteins, as well as those involved in protein transport and secretion (COG-U) exhibited less structural homology between bacteria and archaea (Figures 3D and 3E), deepened the previous findings that sequences of secreted proteins in bacteria evolve fast^63^. Taken together, we observed structural homologies across deep-sea and non-deep-sea proteins, suggesting that DSFC can facilitate the discovery of deep-sea protein with known homologous structures but derived sequences.

In spite of the extensive structural conservation of deep-sea proteins, we still observed structural novelty at the domain level. Using foldseek^64^, we clustered 1.1 million high-quality predicted structures into 61,933 clusters and dissected them into 41,224 high and medium quality domains with the TED Consensus Tool^65^. Exhaustive alignments against the 2.3 million representative structures in AFDB^26, 61^ predicted 392 novel domains with no structural homologs (Figures 3F and 3G). Among those, 25% (98/392) featured extruded β-solenoid structures (Figure 3G), while some small domains exhibited lower symmetry and compact architectures. Notably, 58 symmetrical domains were detected via SymD^66^ (Figure 3H). Thus, these results suggest that the deep-sea harbors untapped structural diversity. Continued exploration of deep-sea metagenomes, therefore, likely will shed light on the dark matter of the protein structure universe.

### Structure-aided deep-sea bioprospecting

Deep-sea environments have driven the evolution of microbial genomes^67, 68^, yielding unique genetic resources for the discovery of functional enzymes such as methyltransferases and helicases (Figure 2H). Helicases, as motor proteins that control sequencing speed in nanopore sequencing^69–71^, represent a promising bioprospecting target for enhancing sequencing throughput. Thus, here, we focus on the helicase superfamily 1 from the DSGC as a test case to demonstrate the biotechnological potential of novel genes from the deep-sea. By leveraging the structural homologies of proteins, we developed a pipeline to mine deep-sea helicases from DSGC via hierarchical alignments with reference structures from three superfamily 1 helicase families (UvrD/Rep, Pif1-like, Upf1-like)^72, 73^ (Figure 4A). Initial alignments (query coverage > 0.75, TM-score > 0.5) were conducted with representatives in DSFC against reference structures to identify candidate gene clusters. To address potential intra-cluster structural heterogeneity, structures for cluster members were predicted using a DSGC-optimized AlphaFold2 (Methods, Figures S4A–C), with structure alignments excluding 46% (1,124/2,432) discordant members (query coverage < 0.75 or TM-score < 0.5). After filtering out sequences with homologs in the *nr* database^74^, OM-RGC or TSGC, this yielded 1,308 DSGC-exclusive helicase candidates exhibiting structural conservation and uncharacterized sequences (Figure 4A). Supporting our structure-guided approach, 78% (1,020/1,308) of these helicase candidates were not identified as such based on the EggNOG database^75^, corroborating issues with sequence-based annotations of these deep-sea genes due to their distinct sequences.

**Figure 4.**
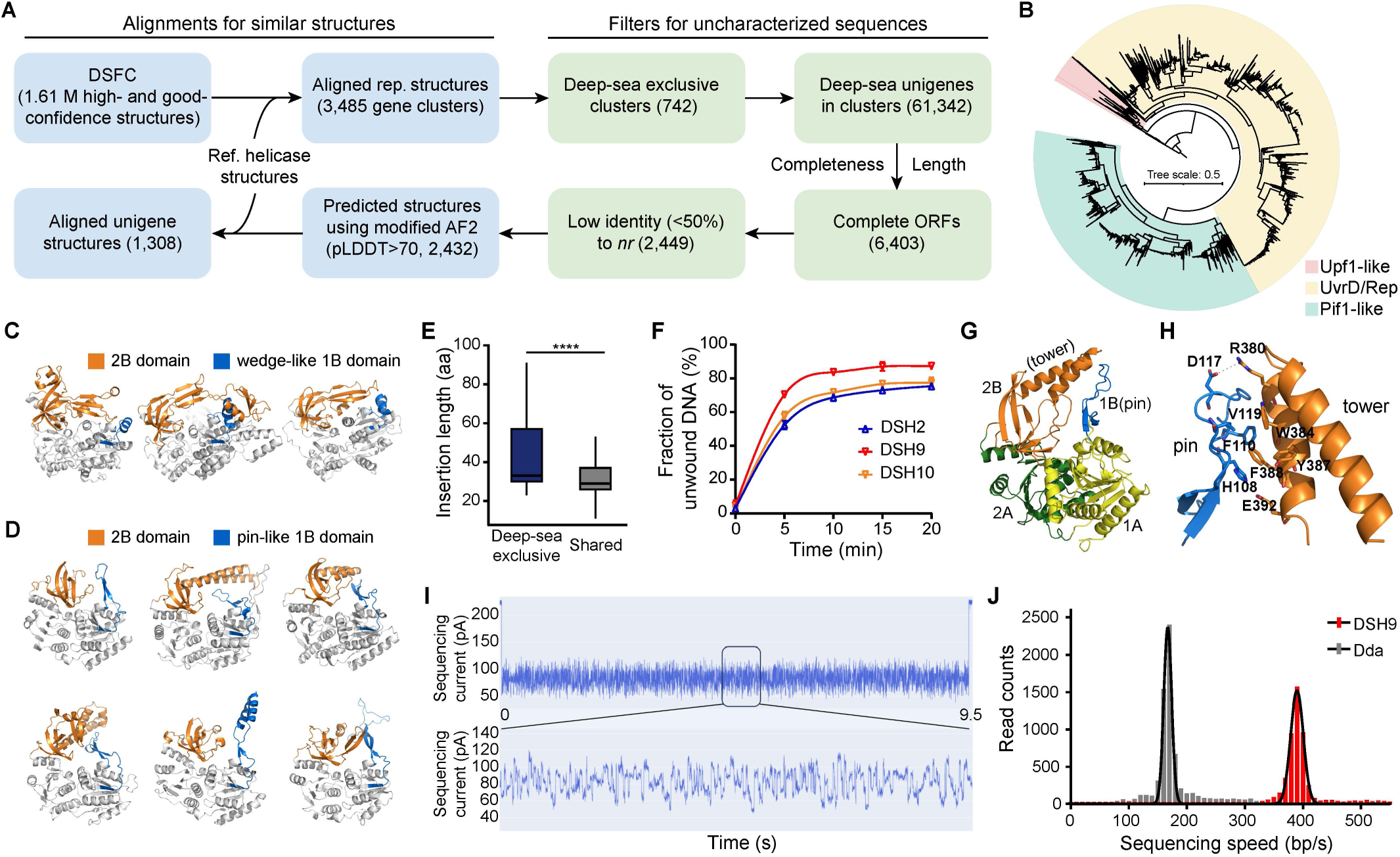
Structure-based mining of a deep-sea helicase with faster sequencing speed. **(A)** Pipeline to mine helicases from DSGC-DSFC datasets based on structural similarity. Deep-sea exclusive clusters refer to clusters without homologs in either the upper ocean or topsoil when clustering at > 20% amino acid identity. **(B)** Tree view of clustering results of the 1,308 mined helicases from DSGC-DSFC. Clustering analysis was performed using the UPGMA algorithm based on TM-score with one superfamily 2 helicase (PDB:1OYW) as outgroup. Shadows indicate the helicase family (UvrD/Rep, Pif1-like, or Upf1-like) assigned to each deep-sea candidate based on structural alignment. **(C, D),** Examples of Pif1-like deep-sea candidates with wedge-like 1B domains (**C**) or pin-like 1B domains (**D**). **(E)** Insertion lengths of 1B domain of deep-sea superfamily 1 helicase candidates (belongs to deep-sea exclusive clusters) and shared ones. Significances were calculated using a two-tailed Wilcoxon test (\*\*\*\**P* < 0.0001). **(F)** The dsDNA unwinding kinetic curves for DSH2, DSH9, and DSH10, respectively. **(G)** The whole structure of apo-DSH9. Four canonical domains were colored as follows: 1A (yellow), 2A (green), 1B (blue), and 2B (orange). **(H)** Key interactions between the pin and the tower in DSH9. **(I)** Nanopore sequencing current trace of an ssDNA measured using DSH9. **(J)** Distribution of sequencing speed using DSH9 and Dda. The black lines represent the nonlinear fitting results.

Structure-based clustering confirmed family-specific groupings of candidate genes (Figure 4B). Among these unique deep-sea helicase candidates, we identified structural diversity especially in Pif1-like helicases concerning their 1B and 2B domains (Figures 4C and 4D). Approximately 18.83% of these structures exhibited a wedge-like 1B domain consisting of one or two α-helices (Figure 4C), which was relatively simple and compact, and may be linked to their specialized role in unwinding G-quadruplex (G4) structures *in-vivo*^76^. In contrast, the 1B domains of other Pif1-like candidates exhibited pin-like structures, which were characterized by extended β-hairpins with variable inserted structures (Figure 4D). Notably, the 1B domain of the deep-sea helicases was significantly longer than their conserved homologs not unique to the deep sea (Figure 4E). Similarly, the 2B domain of these Pif1-like helicase candidates exhibits variations of structural insertions (Figures 4C and 4D).

To assess the functional role of the structural variations in the 1B and 2B domains, we performed experiments on ten selected deep-sea Pif1-like helicase candidates (denoted as DSH1–DSH10) with different 1B sequences (Figure S4D). Three proteins (DSH2, DSH9, and DSH10) were successfully purified through ssDNA cellulose affinity chromatography, demonstrating their ssDNA binding ability (Figure S4E). All three proteins exhibited robust 5’–3’ dsDNA unwinding activity with efficiencies ranging from 70% to 90% (Figure 4F), corroborating their classification as superfamily 1B helicases^73^. In addition, a 2.1-angstrom (Å) resolution X-ray structure of apo-DSH9 revealed similar canonical domains (1A, 2A, 1B, and 2B) with known superfamily 1 helicases but distinctive structural features (Figure 4G and Table S4). Notably, compared to RecD2^77, 78^ and Dda^79^, DSH9 had a longer loop within its pin domain, which extends further alongside the two helices of the tower toward its end (Figure 4G). This configuration creates a larger pin-tower interaction interface (365 Å^2^ vs. Dda’s 208 Å^2^), which is stabilized by hydrophobic contacts (His108/Phe110/Val119–Trp384/Tyr387/Phe388) and hydrogen bonds (His108–Glu392) absent in Dda (Figures 4H and S4F). These interactions likely enhance the mechanical coupling between ATP hydrolysis and DNA unwinding^79^, which therefore may benefit nanopore sequencing.

In nanopore sequencing experiments, we used DSH9 to achieve continuous sequencing reads with clear and consistent squiggling signals and no significant stalling (Figure 4I). Quantitative analysis of 5,000 reads revealed a sequencing speed of 390±11 bp/s, over twice as fast as Dda (167±8 bp/s), with minimal speed variability (Figure 4J). The narrow distribution of DSH9’s sequencing speeds further indicates strong uniformity across different reads. Thus, DSH9 offers advantages in sequencing speed and shows potential for nanopore sequencing.

In summary, we exploited the biotechnological potential of DSGC and DSFC by successfully mining helicases with favorable structural characteristics and sequencing property as a proof of principle for structure-guided deep-sea bioprospecting.

## Discussion

Since the mid-20th century, but particularly during the last two decades, pioneering explorations and technological advancements in deep-sea research have provided a step-change in our understanding of the largest but most unknown ecosystem on Earth. By analyzing metagenomes generated from newly collected samples during the MEER project as well as published deep-sea metagenomes, we studied the global deep-sea microbiome from individual sequences to their potential for biotechnological applications, which is unprecedented. Due to comprehensive sampling and sequencing carried out during the MEER project^13^, the DSGC increases the gene diversity of the Global Ocean Microbiome Catalog^32^ by 51%, thus, significantly extending our knowledge of global marine genetic diversity. Furthermore, our new datasets contribute over 217 million non-redundant genes obtained from hadal zone ecosystems, the most difficult to access deep-sea habitats because of depths between 6 and 11 km and their V-shaped habitat depressions. This resource represents a fundamental step-change, transforming the deep-sea microbiome from a data-sparse frontier into a high-resolution map of planetary genetic diversity.

Our study reframes the deep sea not just as a reservoir of biodiversity, but as a unique evolutionary engine driving the emergence of functionally distinctive enzymatic traits. Deep-sea environments, with their physical isolation and extreme physicochemical conditions, such as high hydrostatic pressure and distinct geochemical regimes, serve as hotspots for evolution^4, 12, 14^. Our convergent approach, linking rapid evolution with ecological prevalence, provides a robust means to navigate this complexity. It implies that the deep-sea environment exerts stringent selective pressures that directly favor certain functional innovations (e.g., thermo-adaptation), while also tolerating or indirectly promoting the fixation of unique genetic variants. These variants can possess specialized properties with biotechnological potential, regardless of whether those properties were adaptive, incidental, or neutral in situ. This dynamic is substantiated by the isolation of proteins with unique sequences and specialized properties: a thermostable Cas9, whose thermostability is likely an adaptive trait, and a Pif1-like helicase with faster sequencing speed, representing the type of distinctive function that can emerge from deep-sea evolution. Consequently, our work positions the deep sea as a critical frontier for evolutionary bioprospecting, where unique environmental conditions act as a filter and catalyst, generating proteins with distinctive properties that are rare in surface ocean and land ecosystems.

Furthermore, our study establishes a scalable blueprint for culture-independent bioprospecting, overcoming the historic bottleneck of relying on isolated microbes. Structure-based approaches have been widely adopted to uncover new enzymes, paving the way for advancing biotechnology^80–82^. The interrogation of our deep-sea datasets (DSGC-DSFC) using the latest structure-analysis tools^64, 83^ in combination with biochemical analyses facilitates the discovery of functional novelty in the deep sea, which includes those from genome editing to nanopore sequencing. Additionally, the effective selection of candidate proteins from DSFC and the usage of DSGC-optimized-AlphaFold2 pronouncedly reduced the computational cost, which makes such an approach more feasible and affordable^84^. Moreover, DSGC-DSFC datasets supply unexplored novelty to the protein sequence and structural space^85^, possibly shedding light on longstanding evolutionary questions including the origin of life on Earth, which is assumed to have taken place in the deep sea^86^.Our work also implies that exploitation of metagenomic resources from extreme environments such as the deep sea as an example is crucial to obtain a more complete assessment of the genetic diversity of our biosphere. At the same time, the novel gene sequences uncovered here provide valuable data to advance AI-driven biological discovery, such as protein structure prediction^87, 88^ and multimodal generative language models^89, 90^.

## Methods

### Sample collection

Published deep-sea metagenomic sequencing data were obtained from NCBI (https://www.ncbi.nlm.nih.gov/). On the basis of information from either the related papers or NCBI metadata, samples were filtered if they were collected in the ocean area deeper than 1,000 m and sequenced with metagenomic pair-end libraries. Finally, 944 deep-sea metagenomic datasets were downloaded for analysis (Table S1). Then we categorized the 944 samples into four ecosystem groups based on the sampling information (Table S1). There were 305 samples assigned to the group of hydrothermal vent ecosystems, and 167, 66, and 406 for the groups of methane seep ecosystems, hadal ecosystems (> 6,000 m in depth), and the other deep-sea ecosystems, respectively. As the published samples collected from the hadal ecosystems were scarce, we additionally integrated 1,194 sediment samples generated by the Mariana Trench Environment and Ecology Research (MEER) project^13^ (Table S1). The samples from the MEER project were assigned to the group of hadal ecosystems. The methods of metagenomic DNA extraction and sequencing of the MEER sediment samples were described in Xiao et al. 2025^14^. Briefly, microbial DNA was extracted from 0.5 g thawed sediment samples using MGIEasy Environmental Microbiome DNA Extraction Kit (MGI). Libraries were prepared with MGIEasy Fast FS DNA Library Prep Set (MGI), followed by a 150 bp paired-end sequencing on DNBSEQ-T series platform.

### Construction of DSGC

All the raw reads were quality-controlled using Fastp (v0.23.1)^91^ with default parameters (-f 0 -t 0 -F 0 -T 0 -Y 30 -l 15) to remove the low-quality, adapter contaminated and duplicated reads. The filtered reads were assembled with MEGAHIT (v1.2.9)^92^ with the option of --presets meta-sensitive. For the 944 published datasets, the threshold for contig filtering was 500 bp. Considering the high sequencing depth of the other 1,194 MEER samples, the contigs shorter than 1,000 bp were removed from the assemblies. According to the assembly results, MetaGeneMark (v3.38)^93^ was used to predict the coding sequences (CDSs) longer than 100 bp in the assembled contigs, and the CDSs were translated into protein sequences according to codon table 11^94^. In total, 1.90 billion protein-coding genes were obtained, which completeness were determined based on the presences of both start and stop codons. All sequences were clustered using MMseqs2 (v14-7e284)^95^ with the following options: easy-cluster --min- seq-id 0.95 -c 0.90 --cov-mode 1 --cluster-mode 3 (set other parameters as default), resulting in a nonredundant microbial gene catalog (namely DSGC) comprising 501,998,347 genes.

In addition, to generate metagenome-assembled genomes (MAGs), MetaWRAP (v1.3.2)^96^ was employed for binning of all the 2,138 deep-sea metagenomes with default parameters. All MAGs were evaluated using CheckM (v1.0.12)^97^, and those with low-quality (completeness <50% or contamination >10%) were removed. All the retained MAGs were clustered at 95% ANI with dRep (v3.3.0)^98^. GTDB-TK (v2.3.2)^99^ was used for taxonomic annotation of each MAG by searching the GTDB (r214)^100^.

### Validation and annotation of DSGC

To calculate the coverage of DSGC to deep-sea microbial gene contents, we randomly selected 100 metagenomic samples from each of the four deep-sea ecosystem groups and extracted up to 50 M reads for each sample. The reads for each sample were aligned to DSGC using DIAMOND (v2.1.6)^101^ with parameters: --outfmt 6 -k 1 --id 70 --query-cover 80 -p 40 -e 1E-04 --faster, and the reads mapping rate of each metagenomic dataset was calculated. The use of protein-level alignment was necessitated by the scale of the DSGC (∼300 Gb of all CDS sequences), providing a balance between computational efficiency and sensitive homology search. The protein sequences in DSGC were clustered at > 20% amino-acid identity with MMseqs2 (v14-7e284)^95^ easy-cluster (--cluster-mode 0 -c 0.5 --min-seq-id 0.2 --cov-mode 2, and other parameters left with their default values) to assign incomplete unigenes to validate the incomplete unigenes based on their homology to complete genes.

For taxonomic annotation of DSGC, genes from all samples were aligned against the Non-Redundant Protein Sequence Database (the *nr* database) (v20200619)^74^. The GTDB-TK taxonomic annotation was assigned to a gene if included in an MAG, and the taxonomic assignment of other genes in DSGC was performed using the lowest common ancestor (LCA) approach based on the aligning results against the *nr* database. We used DIAMOND (v2.1.6, blastp -e 1E-05)^101^ and eggNOG-mapper (v2.0.1, default parameters)^102^ to functionally annotate DSGC with the KEGG database (r104.1)^103^ and the eggNOG database (v5.0.0)^28^, respectively. For the TSGC and OM-RGC datasets, annotations against the eggNOG database (v5.0.0) were carried out using eggNOG-mapper version (v2.0.1) with the same parameters for DSGC (default parameters). Sequences in DSGC, as well as those in global topsoil gene catalog (TSGC^30^, 159.7 M genes) and the *Tara* ocean’s OM-RGC^27^ (46.8 M genes) were aligned against all 21,979 Pfams in the Pfam database (v36.0)^104^ using hmmsearch program (E-value ≤ 0.01) from the HMMER package (v3.3.2)^105^ and then filtered by a sequence coverage of 75%. The Nf (number of effective sequences in MSA) was calculated using plmc (commit 1a9ale9228a2177c618c69040ea8cfc2d902d9df)^106^ with the parameters: --fast -m 1 -n 10. Among the 21,979 Pfams, we detected 18,286 Pfams with at least one homologous from one of the three gene catalogs, and DSGC exhibited the highest Nf in 14,708 Pfams. Protein family-level clustering was performed comparing i) DSGC against the global ocean gene catalog 1.0 (GOGC)^32^, ii) DSGC against both TSGC^30^ and OM-RGC^27^, with MMseqs2 (v14-7e284)^95^ with following the options employed in the GMGC study^33^: easy-cluster --cluster-mode 0 -c 0.5 --min-seq-id 0.2, and we additionally used --cov-mode 0 to cluster sequences with similar length, while other parameters were kept default. As the DSGC, OM-RGC, and TSGC differ in size, we further conducted five independent clustering replicates, each using 10 million sequences randomly subsampled per catalog with the ‘sample’ module of SeqKit (v0.4.5)^107^. All MMseqs2 parameters remained identical to those employed for clustering the full catalogs.

To enable a taxonomically consistent comparison of genetic diversity, all prokaryotic genes in DSGC, OM-RGC, and TSGC were annotated using MMseqs2 (v18.8cc5c)^95^ with the easy-taxonomy routine against the GTDB database (release r226) with “--tax-lineage 1”. As some phyla (or order) were represented by very few genes, a phylum (or order) was considered robust in a given catalog if it comprised >0.1% (or >0.01%, respectively) of the total genes in that catalog. Taxonomic groups that passed this threshold in all three catalogs were defined as overlapping phyla (or orders). DSGC exhibited largest non-redundant gene numbers in these overlapping phyla (or orders) among the three catalogs (Table S3), while it harbored largest non-redundant gene counts in 11/15 overlapping phyla and 45/62 overlapping orders (Table S3).

### Clustering and distribution profiling of COG-L category genes

We calculated the proportion of COG-L genes in microbial MAGs generated from both deep-sea and upper-ocean metagenomes. For the upper-ocean MAGs, we employed the 2,631 MAGs raised by the *Tara* Ocean project^108^. GTDB-TK (v2.3.2)^99^ was used for taxonomic annotation of these MAGs by searching the GTDB (r214)^100^. The coding DNA sequences (CDSs) of all MAGs from deep sea and upper ocean were predicted using Prodigal (v2.6.3)^109^. We used eggNOG-mapper (v2.1.12)^102^ with default parameters to functionally annotate CDSs with the eggNOG database (v5.0.2)^28^. Before comparison, low-quality MAGs with either a completeness < 90% or a contaminant > 5% were removed.

All genes annotated as COG-L category from DSGC, OM-RGC, and TSGC were taxonomically annotated using MMseqs2 (v18.8cc5c)^95^ easy-taxonomy with GTDB (r226) with --tax-lineage 1, and clustered using MMseqs2 (v14-7e284)^95^ with the parameters used in clustering all sequences in the three catalogs: easy-cluster --cluster-mode 0 --cov-mode 0 -c 0.5 --min-seq-id 0.2 (set other parameters as default). Among the 728,098 COG-L category gene clusters, 92.23% showed functional consistency within each cluster, with most of the clustered genes (≥ 90%) were assigned to the same COGs.

To quantify the abundance of all deep-sea COG-L genes, we followed the methods used in the GMGC study^33^, where gene abundance was estimated as the number of short reads mapping to a given sequence. In brief, minimap2 (v2.28-r1209)^110^ was employed to align reads to these genes with default parameters, and the relative abundance of each gene was computed with the “dist1” option of NGLess (v1.5.0)^111^ to distribute multiple mappers (short reads mapping to > 1 unigenes) by unique mapper abundance. The results were normalized by library size to get relative abundance, and the abundance of each gene cluster was determined by summing up the relative abundances of genes in that cluster. The detection number of each gene cluster was the count of samples with non-zero relative abundances.

COG enrichment in exclusively deep-sea cluster (excl-DS-clusters) was conducted using a two-tailed Fisher’s exact test^112^, where deep-sea gene counts in excl-DS-clusters were compared against the background gene counts in shared clusters. Seurat (v4.0.0)^113^ was used to cluster samples with the abundances of either all prevalent (detection > 100) gene clusters or only the prevalent clusters with enriched COG terms. The original Louvain algorithm was used to clustering the samples under the resolution of 0.5, uniform manifold approximation and projection (UMAP) with default parameters (n.neighbors = 30L, min.dist = 0.3) was used for dimensionality reduction. The association between deep-sea ecosystem types and gene distribution was calculated using the Cramer’s *V* test^114^.

### Phylogenetic analysis of the hydrothermal vent-specific DsCas9

The phylogenetic tree was constructed based on amino acid sequences of proteins in a hydrothermal vent-specific Cas9 cluster, two thermostable Cas9s (thermoCas9^42^ and IgnaviCas9^43^) and all well-annotated prokaryotic Cas9s in Swiss-Prot^115^ (UniProt accession numbers: A0Q5Y3, A1IQ68, C9X1G5, G3ECR1, J3F2B0, J7RUA5, Q03JI6, Q03LF7, Q0P897, Q6NKI3, Q8DTE3, Q927P4, Q99ZW2, Q9CLT2). MAFFT (v7.310)^116^ was used to align the sequences, while IQ-Tree (v3.0.1)^117^ was employed to generate the maximum-likelihood tree, and both analyzes were performed with default parameters.

### Constructions of DsCas9 expression vectors

The *E. coli* codon-optimized sequence of *DsCas9* was synthesized (GCATbio) and cloned into the pET28a(+) vector with a C-terminal hexa-histidine tag. Both non-coding and CRISPR-array regions of *CRISPR-DsCas9* locus were also synthesized (GCATbio) and cloned into pCYC184. For small RNA sequencing, the pET28a vector containing the effect protein (pET28a(+)-*DsCas9*) and the pCYC184 vector containing the noncoding RNA (pCYC184-*DsCas9nc*) were co-transformed into BL21(DE3) competent cells for heterologous expression of the CRISPR-DsCas9 system in bacteria. For human genome editing assay, the human codon-optimized sequence of *DsCas9* with bpNLS tags and a U6-driven sgRNA expressing cassette with a *Bsa*I golden gate site were synthesized (GenScript Life Science) and cloned into a modified PX458 vector, resulting in a plasmid named as PX458-*DsCas9*. The target oligos were synthesized, annealed, and inserted into the *Bsa*Ⅰ site of PX458-*DsCas9* to generate target editing plasmids for human genome editing.

### Small RNA sequencing and data analysis of CRISPR-DsCas9

For small RNA sequencing, the *E. coli* carrying the heterologous *CRISPR-DsCas9* system was cultured and harvest for RNA extraction. The total RNA was extracted from 5 mL of cell culture using the RNeasy PowerFecal Pro Kit (QIAGE) according to the manufacturer’s protocol. Subsequently, the small RNA sequencing library was prepared using the by MGIEasy Small RNA Library kit (MGI) according to the manufacturer’s protocol and sequenced on the DNBSEQ-T7 sequencer (MGI) with paired-end 150 bp (PE150) reads. For data analysis, the obtained sequencing reads were aligned to the pCYC184-*DsCas9nc* plasmid using the BWA (v0.7.1)^118^ MEM algorithm with default parameters. The resulting SAM files were converted to BAM format, sorted, and indexed using SAMtools (v1.20)^119^. Uniquely mapped reads were extracted and analyzed using SAMtools to evaluate sequencing depth, coverage, and read distribution across the plasmid. To determine the mature crRNA and tracrRNA sequences, mapped reads flanking the CRISPR locus were analyzed. Read coverage around the CRISPR array was examined to detect distinct peaks corresponding to processed crRNA and tracrRNA. The mapped reads and coverage profiles were visualized using Integrative Genomics Viewer (IGV) (v2.18.4)^120^ to identify the potential processing patterns of crRNA and tracrRNA.

### Protein expression and purification

The *E. coli* BL21(DE3) cells containing the pET28a-*DsCas9* plasmid was incubated at 37°C, 200 rpm until the optical density at 600 nm (OD600) reached 0.6, and then induced with 0.5 mM IPTG overnight at 16°C, 200 rpm for protein expression. The cells were collected by centrifugation and resuspended in lysis buffer (20 mM Tris-HCl pH 7.5, 1M NaCl, 20 mM imidazole, 1 mM PMSF). The cells were disrupted by sonication, and the lysate was clarified by centrifugation. The clear lysate was loaded onto a Ni-NTA gravity column (HisSep Ni-NTA Agarose Resin, Yeasen Biotechnology) and incubated for 1 hour. The column was washed with 10–15 column volumes of lysis buffer, and the proteins were eluted with 1–2 column volumes of elute buffer (20 mM Tris-HCl pH 7.5, 300 mM Imidazole, 1M NaCl). The eluted proteins were diluted with dilution buffer (20 mM Tris-HCl pH 7.5, 1 mM DTT, 10% glycerol) and further purified on a Heparin column (HiTrap Heparin HP, Cytiva) for affinity chromatography purification (ӒKTA pure 25, Cytiva) with a linear gradient of 0.1–1M NaCl. The purified proteins were concentrated and further fractionated by size-exclusion chromatography (SEC) using a Superdex 200 Increase 10/300 GL column (Cytiva). The purified proteins were concentrated to 1–5 mg/mL and stored at –80°C in storage buffer (50 mM Tris-HCl, 250 mM NaCl, 1 mM DTT, 30% glycerol).

### Protein thermal stability assay

To compare the thermal stability of SpCas9 and DsCas9, we conducted melt assays using a protein thermal shift dye kit (ThermoFisher Scientific) in a real-time PCR instrument (ABI) following the manufacturer’s instruction. Briefly, the protein melt reaction, containing approximately 1 μg sample protein, protein thermal shift dye, and protein thermal shift buffer, was prepared on ice with three replications. The melting temperatures (Tm) of the proteins were calculated based on the melt curves obtained after performing the experiments in accordance with the protein thermal shift dye kit protocol.

### In-vitro dsDNA cleavage assay

The guide RNAs with targets of SpCas9 and DsCas9 were obtained by *in-vitro* transcription using the MEGAshortscript kit (ThermoFisher Scientific) and purified with the Guide-it IVT RNA Clean-Up Kit following the manual protocol. The dsDNA substrates for SpCas9 and DsCas9 were prepared by PCR with the same target sequence and specific PAM sequence. To compare the *in-vitro* dsDNA cleavage activity of SpCas9 and DsCas9 at various temperatures, a mixture containing 25 nM Cas protein, 100 nM gRNA, and 1×NEBuffer3.1 was prepared and incubated at tested temperature for 10 minutes. Subsequently, 100 ng of plasmids or PCR products were added, and the incubation continued at the same temperature for 20 minutes. The reaction was terminated with 2.5 mM EDTA and digested with protease K at 55°C for 10 minutes, followed by loading for DNA gel electrophoresis. To determine the effect of salt concentration on the *in-vitro* dsDNA cleavage activity of SpCas9 and DsCas9, a mixture containing 25 nM Cas protein, 100 nM gRNA, and 1×Basic Reaction Buffer (50 mM Tris-HCl, 10 mM MgCl2, 100 μg/mL BSA, pH 7.9) with a salt concentration gradient was prepared and incubated at 37°C for 10 minutes. Then, the *in-vitro* dsDNA cleavage was performed as previously described and analyzed with DNA gel electrophoresis.

### Cell culture, transfection and genome editing assay

The HEK293T cells were cultivated with DMEM medium supplied with 10% fetal bovine serum (Gibco). The cells were initially seeded in a 12-well plate at a density of 100,000 cells per well one day before transfection. Plasmid transfection was performed with Lipo8000 (Beyotime Biotechnology) according to the manufacturer’s protocol. The transfection efficiency was assessed, and the cells were harvested for evaluation 2 to 4 days after transfection. The genome DNA was extracted using the TIANamp Genomic DNA Kit (TIANGEN). The genome editing activity was detected by next-generation sequencing with a PCR amplicon library (Novogene). The genome editing efficiency was calculated by CRISPRessoPooled (v2.3.3) software^121^ (https://github.com/pinellolab/CRISPResso2).

### Selection test for shared protein clusters among DSGC, OM-RGC, and TSGC

To investigate the evolution of deep-sea genes, we examined the presence of rapidly evolved genes using a branch-site model, adaptive branch-site random effects likelihood (aBS-REL)^122^. Given the challenge of strict ortholog-paralog distinction in metagenome-derived gene catalogs, we performed this evolutionary analysis by following the homology-based practice as established in the GMGC pipeline^33^. Briefly, the COG-L category gene clusters shared by DSGC, OM-RGC, and TSGC were used for direct comparison. As incomplete ORFs may confound the computation, we only tested complete genes, and clusters with more than three complete members from each of the three gene catalogs and less than 500 complete members in total were selected for this analysis. In addition, we were only focused on clusters that were detected in more than 100 deep-sea samples. Algorithm super5 of muscle (v5.1)^123^ was employed to generate multi-sequence alignments, and the alignments was converted into codon-based DNA alignment using pal2nal (v14)^124^. Phylogenetic tree was constructed using FastTree (v2.1.10)^125^, and phylogenetic diversity of genes from each of the three catalogs was assessed using GenomeTreeTK (v0.1.8, github.com/dparks1134/GenomeTreeTk). Selection tests were run using the aBS-REL mode of HyPhy (v2.5.48)^126^ with parameters of “--branches Leaves”. Finally, we tested 5,837 protein-family clusters containing 1,100,453 genes form DSGC, OM-RGC, and TSGC. Genes were defined as “rapidly evolving” if they exhibited a statistically significant signature (Holm-Bonferroni corrected *P* < 0.05) of episodic diversifying positive selection, as detected by the aBS-REL model^122^ in HyPhy^126^. Additionally, we also tested these selection test results in 100 randomly selected clusters with a slower but more accurate pipeline, where we employed MAFFT (L-INS-I, v7.407) and IQ-Tree3 (v3.0.1) instead of muscle (super5) and FastTree and used TrimAL (v1.5.rev0, - gappyout)^127^ to remove poorly aligned regions in MSA (Figure S2L).

To validate the results of the selection test on deep-sea genes, we tested whether the rapidly evolved genes had higher nucleotide diversity, pN/pS, and detection rates than the others. The nucleotide diversity and pN/pS analysis was limited to 57,013 genes, which were involved in the selection test and associated with the MAGs generated in this study. Bowtie2^128^ was used to align metagenomic reads of related samples against all MAGs clustered at 95% ANI, and InStrain (v1.9.0)^129^ was used to calculate the nucleotide diversity and pN/pS of each gene. The Wilcoxon test was used to investigate if rapidly evolved genes are more prevalent (with higher frequencies of detection) in deep-sea than other genes, and a false discovery rate correction (*P* < 0.05) was implemented using the Benjamini-Hochberg procedure^130^.

### Construction and validation of DSFC

The protein family-level clusters with more than 30 members from DSGC were included in the construction of DSFC. For each cluster, all sequences predicted as complete ORFs were aligned to the centroid (the representative sequences clustered >20% amino acid identity) using blastp (v2.16.0)^131^, and the top-two sequences hit on centroids were subjected to ESMFold^23^ with default parameters for structure prediction. After prediction, the structure with a higher pTM score in each cluster was chosen as the representative and was deposited in DSFC.

To assess the representativeness of each representative structure across all protein structures in the respective sequence cluster (clustered at ≥ 20% amino acid identity), we randomly selected 220 clusters for benchmark validation. The size of each cluster varied from dozens to hundreds of members, encompassing a total of 13,337 proteins with complete ORFs. All protein structures were predicted using the default parameters of ESMFold^23^, and we generated 10,156 high-confidence structures (pLDDT > 0.7 and pTM > 0.7). Subsequently, the high-confidence structures within each cluster were aligned to the representative structure using TM-align (r2022/4/12)^132^. Structural alignments were filtered by TM-score > 0.5 (normalized by length of query). The representative structure confidently reflects those of most members (> 75%) in the same cluster out of > 91% tested sequence clusters (Figure S3B).

Based on the pLDDT and pTM scores generated by ESMFold, all structures in DSFC were assigned to three groups, e.g. high confidence structures (pLDDT > 0.7 and pTM > 0.7), good confidence structures (pLDDT > 0.5 and pTM > 0.5), and low confidence structures (pLDDT ≤ 0.5 or pTM ≤ 0.5). We randomly selected 1,000 proteins with high-confidence structures in DSFC, each of which is a representative of the 61,933 structural clusters (see *Identification of novel structural domains*), and submitted their sequences to AlphaFold3 online server (AF3) to predict the structures^133^. Then, the average pLDDT scores of the best models generated by AF3 (model_0) were calculated, and high confidence AF3 models (average pLDDT > 0.7) were chosen for pairwise comparison against ESMFold models in TM-score (r2022/2/27)^134^ (Figure S3C) to assess the robustness of our high confidence models generated by ESMFold.

### Structural similarity and functional analysis of DSFC

The script “foldseek.sh” in Nomburg et al. (2024)^58^ with default parameters was used for pairwise structure alignments. Briefly, the search routine of Foldseek (v9.427df8a)^135^ was used to align high-confidence structures in DSFC against 2.3 million protein cluster centroids from the AFDB^61^, all experimental structures in PDB (v20240825)^62^, and themselves. For alignments against AFDB and PDB, the best hits were filtered based on a TM-score > 0.5 and E-value < 0.001. For alignments among structures in DSFC, self-alignments were excluded, and the best hits were filtered by coverages (both query cov and target cov) > 0.9 and TM-score > 0.5.

To evaluate the structural conservation across archaea and bacteria, we performed clustering of the high-confidence structures categorized to Archaea or Bacteria using Foldseek (v9.427df8a, easy-cluster -c 0.9 -e 0.01)^135^. TMHMM (v2.0c)^136^ and SignalP (v6.0)^137^ were used to predict transmembrane domains and signal peptides, respectively. Proteins were classified as follows: if a protein exhibited a secretion probability above 0.75 and contained no transmembrane domains, or contained one transmembrane domain within the first 60 residues in sequence, it was assigned as a secretory protein; proteins lacking both transmembrane domains and signal peptides were classified as cytoplasmatic, while all others were assigned as transmembrane proteins. Enrichment analysis of COG categories was calculated using the Fisher’s exact test^112^. For a certain COG category, numbers of structures within the clusters exhibited cross-domain homologies were compared against the numbers of other structures.

### Identification of novel structural domains

We used Foldseek easy-cluster (v9.427df8a)^135^ to cluster 1,119,421 high-quality predicted structures into 61,933 clusters (-c 0.7 --cov-mode 0 -e 0.01). The representative structures of all clusters were parsed into 192,890 domains by the TED consensus tool (v1.0)^65^, which integrates three domain parsers—Merizo^138^, Chainsaw^139^ and UniDoc^140^—where domains identified by all three or any two tools were defined as high and medium quality domains, respectively, totaling 104,508. Following the TED workflow, we applied a series of quality filtering criteria, including packing density > 10.3, pLDDT80 > 90 and at least 6 secondary structural fragments, to refine the selection to 41,224 domains using the script Fig3F.novel_domain_analysis.sh on our GitHub repository. These filtered domains were then exhaustively searched against 2.3 million representative structures of AFDB clusters using Foldseek easy-search^135^ (-c 0.6 --cov-mode 2 --tmscore-threshold 0.56 --alignment-type 1 --exhaustive-search), resulting in 392 nonhit domains^65^. When we used Foldseek easy-search, the coverage mode was set to 2 to ensure at least 60% of the query deep-sea domain was aligned to the target AFDB whole structure; and the TM-score threshold was set to 0.56 to be in line with the TED protocol^65^ to compensate for the foldseek approximating the true TM-align score. These nonhit domains were considered as relatively novel compared to AFDB. The novelty scores of domains were computed by Foldclass and the domains were segmented by the TED consensus tool^65, 141^. Then, SymD (v1.61)^66^ was used to identify 58 symmetrical domains (Z-score at best alignment > 9).

### Performance assessment of DSGC-optimized AlphaFold2

Following the instruction provided by the official GitHub repository of AlphaFold (https://github.com/google-deepmind/alphafold), we first ran AlphaFold2 (AF2, v2.3) with the default settings, which would search against its default genetic datasets (including BFD ∼1.8 TB, MGnify ∼ 120 GB, UniRef90 ∼ 67 GB, UniClust30 ∼ 206 GB). This would generate an output directory, in which there was a subfolder, called *msas*, containing 3 MSA files, namely bfd_uniref_hits.a3m, mgnify_hits.sto, and uniref90_hits.sto. These three MSA files would later be used to extract the MSA features by AF2 and thus were referred to as the AF2-MSA by us.

We used JackHMMER^142^, which is implemented in HMMER v3.3.2^105^, to search against the deep-sea sequences with the same chosen parameters of AF2. This would generate an MSA file, named deepsea_hits.sto, which was referred to as the DS-MSA by us.

Then, we removed the 3 MSA files in the output folder (saving all output files in the first run somewhere else) and instead renamed the deepsea_hits.sto as mgnify_hits.sto. In this way, as we ran AF2 again and turned on the use_precomputed_msas flag, AF2 would recognize only the substituted STO file as the precomputed MSA and conducted the subsequent inference.

To validate the improvement of DSGC on structure prediction of deep-sea proteins, we randomly selected 200 protein sequences from DSGC, which lengths were equally distributed in the range of 0–1000 residues. Then, we built DS-MSA and AF2-MSA by searching DSGC and the default genetic datasets of AF2, respectively, and performed structure prediction using AF2 with each MSA. The computing time consumed for prediction and the resultant pLDDT score for each query were compared across MSAs (Figures S4A and S4B). The same set of AF2 parameters was used throughout the AF2 prediction with different datasets to ensure consistency and comparability.

To further validate the accuracy of prediction using DSGC optimized AF2, we calculated the structure similarity (represented by TM-score) between the models predicted using DS-MSA and AF2-MSA respectively. The results showed high consistency between these models, especially those with higher pLDDT scores (Figure S4C). Together with Figure S4A and Figure S4B, these results indicated that DSGC optimized AF2’s performance and exhibited high efficiency in predicting structures of a deep-sea protein, while maintaining comparable accuracy to default AF2 predictions.

### Structure-based mining of superfamily 1 helicase

Entries of superfamily 1 helicases with experimental 3D structures in Swiss-Prot^115^ website (Nov. 2024) were filtered manually, and 12 structures belonging to the three families of superfamily 1 helicase were selected as references for helicase mining (PDB accession number: Pif1-like family: 3E1S, 5FHH, 5O6B, 7OTJ; Upf1-like family: 2GJK, 2XZL, 4B3F, 5EAN, 5MZN; UvrD/Rep family: 1PJR, 1UAA, 3LFU)^72, 73^. We then employed Foldseek (v9.427df8a)^135^ to align all structures in DSFC except those with low confidence against these reference structures using the default settings of the script in Nomburg et al. (2024)^58^, and 3,485 structures, which represented 3,485 sequence clusters, were confidently (query coverage > 0.75, TM-score > 0.5) aligned to the reference structures. 742 of these 3,485 clusters were found to be deep-sea exclusive, and we found 6,403 complete ORFs with a protein length in the range of 350–1100 aa (determined by the lengths of the references). These unigenes were then aligned to the nr database (v20200619)^74^, and 2,449 sequences with identity < 50% were kept in total. The structures of the 2,449 sequences were predicted using DSGC-optimized AF2, and structures with average pLDDT > 70 were aligned against the 12 reference structures again with the same threshold (query coverage > 0.75, TM-score > 0.5) to ensure robustness.

Structures of the 1,308 superfamily 1 helicase candidates and the 12 references, and one structure of a superfamily 2 helicase as the outgroup (PDB accession number: 1OYW) were pairwise aligned using US-align (v20240730)^143^, and clustered using the unweighted pair group method with arithmetic mean (UPGMA) method based on pairwise TM-scores.

To compare the insertion length of the 1B domain, we retrieved deep-sea unigenes in clusters that were shared among DSGC, OM-RGC and TSGC (not exclusive to DSGC). These unigenes were filtered based on completeness and protein length (in the range of 350–1100 aa), and their structures were predicted using ESMFold^23^ and aligned against the 12 reference structures. All proteins in both exclusively deep-sea clusters and the shared clusters, which were best aligned to one of the Pif1-like references (PDB accession number: 3E1S, 5FHH, 5O6B, 7OTJ), were used to construct a multiple sequence alignment using the default parameters of MAFFT (v7.407)^116^. The lengths of the 1B domain insertions, which were bounded by the conserved motifs Ⅰc and Ⅱ sequences of the 1A domains, were calculated based on the MSA results.

### Cloning, expression and purification of helicases

The DNA sequences of the full-length deep-sea helicases (DSHs) were ligated into the *Nde*I and *Xho*I sites of the pET-28a(+) plasmid (Fisher Scientific), respectively. Therefore, the N-termini of the expressed DSH proteins have 6×His tag and a thrombin protease cleavage site. The cloned pET-28a(+)-DSH plasmids were transformed into *Escherichia coli* BL21 (DE3) competent cells (Vazyme). Cells were cultured in LB medium with shaking at 37°C until OD600 = 0.6–0.8. After cooling down to 16°C, isopropyl-β-d-thiogalactoside (IPTG) was added at a final concentration of 500 μM to induce protein expression overnight.

The cells expressing DSHs were collected and resuspended with buffer A (20 mM Tris-HCl pH 7.5, 250 mM NaCl, 20 mM Imidazole) before being disrupted. The disrupted cells were centrifuged, after which the supernatant was collected. The supernatant was loaded onto a Ni^2+^ chelation column (Cytiva) and washed with buffer A. Then the target proteins were eluted with buffer B (20 mM Tris-HCl pH 7.5, 250 mM NaCl, 300 mM Imidazole). The eluted proteins were passed through a desalting column (Cytiva) equilibrated with buffer C (20 mM Tris-HCl pH 7.5, 80 mM NaCl) for buffer replacement. The DSH proteins and thrombin protease were then added to buffer C balanced single-stranded DNA (ssDNA) cellulose column (Sigma-Aldrich), incubated at 4°C overnight. We collected the ssDNA cellulose resin and washed it with buffer C. And then the target proteins were eluted with buffer D (20 mM Tris-HCl pH 7.5, 1000 mM NaCl). The proteins purified by ssDNA cellulose column were concentrated and then loaded onto Superdex 200 (Cytiva). The size exclusion chromatography (SEC) buffer was buffer E (20 mM Tris-HCl pH 7.5, 200 mM NaCl). Finally, the target proteins were collected and concentrated to 10 mg/ml, then stored at –80°C before use. The Dda protein was purified as previously described^144^.

### dsDNA unwinding assay of DSH Proteins

ssDNA-1 and ssDNA-2 were annealed into 5’ overhang dsDNA (ovDNA). 0.05 μM ovDNA and 0.1 μM DSH proteins were mixed into the reaction buffer G (100 mM HEPES-K pH 8.0, 0.1 mg/mL BSA, 10 mM MgCl2 and 150 mM KCl). The reactions were initiated by mixing up 1 mM ATP and 1 μM competing ssDNA-3. A microplate reader (Synergy H1, BioTek) was used to detect the kinetic changes of the fluorescence intensity (*I*) within 20 min of the reactions at 30 °C, with three replicates for each group. The positive control (*Ipc*) is the ovDNA replaced with ssDNA-2, while the negative control (*Inc*) was without the addition of DSH proteins. Fraction of unwound DNA = 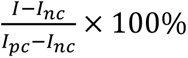.

### Crystallization and crystallography

The apo-DSH9 protein (10 mg/ml in buffer E) was crystallized using the hanging-drop vapor diffusion method equilibrated against a reservoir solution of 200 mM potassium sodium tartrate tetrahydrate, 20% PEG 3350. Crystals were grown to the optimal size for one week at 20°C and frozen in cryoprotectant consisting of the reservoir solution supplement with 30% glycerol.

X-ray diffraction data were collected at the beamline BL18U1 of Shanghai Synchrotron Radiation Facility (SSRF) at a wavelength of 0.9790 Å at – 173°C and processed using the automatic XIA2 software package^145^. The initial structure of DSH9 was determined by the PHASER program^146^ using AF2 predicted model as a searching template. The structure refinement was performed with the PHENIX program^147^, and the model was created by COOT^148^. Detailed data collection and refinement statistics are summarized in Table S4. Ramachandran plot statistics for the apo-DSH9 structure are as follows: 97.31% of residues are in the most favored regions, 2.69% in additionally allowed regions, and without any residues in generously allowed or disallowed regions.

### Nanopore sequencing experiments

All sequencing experiments were performed using the CycloneSEQ WT02 nanopore sequencing platform^149^. Unless stated otherwise, components used for library preparation and sequencing are provided in the CycloneSEQ Universal Library Prep Set and CycloneSEQ WT Sequencing Kit. Briefly, a 3.6-kb dsDNA fragment was first ligated with sequencing adapter, followed by loading of helicases DSH9 or Dda to prepare the sequencing library. Here, we engineered DSH9-V119C/L369C and Dda-E94C/A360C mutants to introduce a pin-tower disulfide bridge, thereby preventing helicase dissociation and improving sequencing continuity. Then, the library was mixed with the sequencing buffer and added to WT Sequencing Flow Cell to perform nanopore sequencing at 30°C. For comparison, data of 5,000 sequencing reads was randomly selected from the sequencing results with either DSH9 or Dda. Sequencing speed profiles were calculated by dividing the DNA fragment length by the corresponding read duration, followed by statistical characterization of the speed distribution patterns. Our comparison intentionally focuses on baseline, non-optimized conditions and thus should be interpreted as a comparison of intrinsic unwinding performance rather than maximum achievable speed.

### Quantification and Statistical Analysis

Data are shown as mean/median ± standard deviation, or, as box plot in which the center line represents the median and the box’s bounds indicate the upper and lower quantiles. The significance differences between two groups were determined using the two-tailed Wilcoxon test, with *P* < 0.05 considered statistically significant. Enrichments were calculated using Fisher’s exact test, and association between clustered samples and their habitat were indicated using Cramer’s value. Exact values of repeat numbers are shown as n in the method details and figure/table legends. The melt curve assay of DsCas9s and dsDNA unwinding assay of DSH proteins was repeated three times. Graph preparation and statistical analyses were performed using R Studio (https://www.r-project.org).

## Data and code availability

All metagenomic data used in this study have been published before and listed in Supplementary Table 1, and we provided example download commands in our GitHub repository (see code availability). Atomic coordinates and structural factors for the apo-DSH9 (PDB: 8JX6) have been deposited at the Protein Data Bank. The sequences and structures of the DSGC-DSFC dataset have been deposited and indexed at a new website (https://db.genomics.cn/exmode/collections/dsgc), and free online services are provided to easily manipulate and search for remotely homologous deep-sea genes and proteins of their interests. Metagenome-assembled genomes, annotations, and predicted structure domains of our dataset have been deposited at Zenodo (https://zenodo.org/records/18170924).

Original code and intermediate results, as well as a step-by-step bioinformatic analysis protocol are available at Github (https://github.com/lidenghui92/DSGC-DSFC). This includes scripts and intermediate results required for most computational steps, and large intermediate files that are necessary to reproduce the figures have been deposited at Zenodo (https://zenodo.org/records/18170924).

## Acknowledgments

This work was supported by the grant ZDKJ2021028 from the Key Research and Development Program of Hainan Province, the National Key R&D Program of China (2024YFC3406300, 2022YFC2805404), the Technological Innovation Projects of Qingdao West Coast New Area (ZDKC-2022-03), the National Natural Science Foundation of China (No. 32100994, U2544219), the Guangdong Basic and Applied Basic Research Foundation (2025B1515020038), the “Pioneer” and “Leading Goose” R&D Program of Zhejiang (2024C03004), the High-performance Computing Platform of YZBSTCACC, the Marine S&T Fund of Shandong Province for Pilot National Laboratory for Marine Science and Technology (Qingdao) (No.2022QNLM030004), the Project of Sanya Yazhou Bay Science and Technology City, Grant No, SKJC-2024-01-002, the Shenzhen Engineering Laboratory for Molecular Enzymology (DRC-SZ [2018]958), and the Major Science and Technology Research Project of Sanya Yazhou Bay Science and Technology City (grant SKJC-2021-01-001). We sincerely thank the Institute of Deep-sea Science and Engineering, Chinese Academy of Sciences (IDSSE CAS) for supporting this research through the deployment of the research vessels R/V *Tan Suo Yi Hao*. Special acknowledgment is given to the teams operating the HOV *Fen Dou Zhe* and HOV *Shen Hai Yong Shi*, whose cutting-edge capabilities have enabled critical deep-sea observations in this study. We appreciate all the assistance provided by the crews on R/V *Kexue*. We thank the staff scientists at beamline BL18U1 of the Shanghai Synchrotron Radiation Facility for providing technical support and assistance in data collection.

## Author Contributions

Yang Guo, Zongan Wang, D. L., G. F., L. W., H. L. conceived this study; Yang Guo, Zongan Wang, D. L., Z. Z., Z. S., C. Q., B. L., B. H., Yuqian Liu, J. Z., Yue Liu, H. C., K. M., T. Y., J. T., Yulu Zhang, Zhijie Wang, and Xiaokai Xu conducted coding and analysis; L. W., H. L., Z. L., X. S., C. K., L. S., A. J., N. Z., Q. J., Junyi Chen, K. C. carried out the experiments; Yang Guo, D. L., Zongan Wang, L. W., H. L., L. M., M. W., Weishu Zhao, Weijia Zhang, H. Z., Y. W., M. H., Xiang Xiao, J. W., Xun Xu, S. L., Jiayu Chen collected samples and data; Yang Guo, Zongan Wang, D. L., L. W., H. L. wrote the original draft; T. M., Xun Xu, G. F., Weishu Zhao, Xiang Xiao, Y. S., I. S., M. W., X. L., L. M. participated in review and editing; Xun Xu, Y. D., S. L., Yuxiang Li, Yang Guo, F. G., L. M., Yue Zheng, T. Z., C. L., J. W., H. Y., B. W., Wenwei Zhang, Ying Gu supervised this study. Yang Guo, Zongan Wang, D. L., L. W., H. L. contributed equally to this work.

## References

1. Jannasch, H.W., and Taylor, C.D. (1984). Deep-sea microbiology. Annu Rev Microbiol 38, 487–514.

2. Danovaro, R., Snelgrove, P.V., and Tyler, P. (2014). Challenging the paradigms of deep-sea ecology. Trends Ecol Evol 29, 465–475.

3. Jamieson, A.J., Fujii, T., Mayor, D.J., Solan, M., and Priede, I.G. (2010). Hadal trenches: the ecology of the deepest places on Earth. Trends Ecol Evol 25, 190–197.

4. Dick, G.J. (2019). The microbiomes of deep-sea hydrothermal vents: distributed globally, shaped locally. Nature Reviews Microbiology 17, 271–283.

5. Zobell, C.E., and Morita, R.Y. (1957). Barophilic bacteria in some deep sea sediments. J Bacteriol 73, 563–568.

6. Sanders, H.L., and Hessler, R.R. (1969). Ecology of the deep-sea benthos. Science 163, 1419–1424.

7. Corliss, J.B., Dymond, J., Gordon, L.I., Edmond, J.M., von Herzen, R.P., Ballard, R.D., Green, K., Williams, D., Bainbridge, A., Crane, K., et al. (1979). Submarine thermal sprirngs on the galapagos rift. Science 203, 1073–1083.

8. Jorgensen, B.B., and Boetius, A. (2007). Feast and famine--microbial life in the deep-sea bed. Nat Rev Microbiol 5, 770–781.

9. Dick, G.J. (2019). The microbiomes of deep-sea hydrothermal vents: distributed globally, shaped locally. Nat Rev Microbiol 17, 271–283.

10. Santelli, C.M., Orcutt, B.N., Banning, E., Bach, W., Moyer, C.L., Sogin, M.L., Staudigel, H., and Edwards, K.J. (2008). Abundance and diversity of microbial life in ocean crust. Nature 453, 653–656.

11. Case, D.H., Pasulka, A.L., Marlow, J.J., Grupe, B.M., Levin, L.A., and Orphan, V.J. (2015). Methane Seep Carbonates Host Distinct, Diverse, and Dynamic Microbial Assemblages. mBio 6, 10.1128/mbio.01348-01315.

12. Ruff, S.E., Biddle, J.F., Teske, A.P., Knittel, K., Boetius, A., and Ramette, A. (2015). Global dispersion and local diversification of the methane seep microbiome. Proc Natl Acad Sci U S A 112, 4015–4020.

13. Xiao, X., Wang, J., and Ding, K. (2025). MEER: Extraordinary flourishing ecosystem in the deepest ocean. Cell 188, 1175–1177.

14. Xiao, X., Zhao, W., Song, Z., Qi, Q., Wang, B., Zhu, J., Lin, J., Wang, J., Hu, A., Huang, S., et al. (2025). Microbial ecosystems and ecological driving forces in the deepest ocean sediments. Cell 188, 1363–1377.e1369.

15. Chen, J., Jia, Y., Sun, Y., Liu, K., Zhou, C., Liu, C., Li, D., Liu, G., Zhang, C., and Yang, T. (2024). Global marine microbial diversity and its potential in bioprospecting. Nature, 1-9.

16. Cavicchioli, R., Charlton, T., Ertan, H., Omar, S.M., Siddiqui, K., and Williams, T. (2011). Biotechnological uses of enzymes from psychrophiles. Microbial biotechnology 4, 449–460.

17. Thornburg, C.C., Zabriskie, T.M., and McPhail, K.L. (2010). Deep-sea hydrothermal vents: potential hot spots for natural products discovery? Journal of natural products 73, 489–499.

18. Rothschild, L.J., and Mancinelli, R.L. (2001). Life in extreme environments. Nature 409, 1092–1101.

19. Pruski, A.M., and Dixon, D.R. (2003). Toxic vents and DNA damage: first evidence from a naturally contaminated deep-sea environment. Aquatic Toxicology 64, 1–13.

20. Marsic, D., Flaman, J.-M., and Ng, J.D. (2008). New DNA polymerase from the hyperthermophilic marine archaeon Thermococcus thioreducens. Extremophiles 12, 775–788.

21. Mattila, P., Korpela, J., Tenkanen, T., and Pitkanen, K. (1991). Fidelity of DNA synthesis by the Thermococcus litoralis DNA polymerase--an extremely heat stable enzyme with proofreading activity. Nucleic Acids Res 19, 4967–4973.

22. Jumper, J., Evans, R., Pritzel, A., Green, T., Figurnov, M., Ronneberger, O., Tunyasuvunakool, K., Bates, R., Zidek, A., Potapenko, A., et al. (2021). Highly accurate protein structure prediction with AlphaFold. Nature 596, 583–589.

23. Lin, Z., Akin, H., Rao, R., Hie, B., Zhu, Z., Lu, W., Smetanin, N., Verkuil, R., Kabeli, O., Shmueli, Y., et al. (2023). Evolutionary-scale prediction of atomic-level protein structure with a language model. Science 379, 1123–1130.

24. Liang, J., Feng, J.-C., Zhang, S., Cai, Y., Yang, Z., Ni, T., and Yang, H.-Y. (2021). Role of deep-sea equipment in promoting the forefront of studies on life in extreme environments. iScience 24.

25. Barrio-Hernandez, I., Yeo, J., Janes, J., Mirdita, M., Gilchrist, C.L.M., Wein, T., Varadi, M., Velankar, S., Beltrao, P., and Steinegger, M. (2023). Clustering predicted structures at the scale of the known protein universe. Nature 622, 637–645.

26. Varadi, M., Bertoni, D., Magana, P., Paramval, U., Pidruchna, I., Radhakrishnan, M., Tsenkov, M., Nair, S., Mirdita, M., Yeo, J., et al. (2024). AlphaFold Protein Structure Database in 2024: providing structure coverage for over 214 million protein sequences. Nucleic Acids Res 52, D368–D375.

27. Salazar, G., Paoli, L., Alberti, A., Huerta-Cepas, J., Ruscheweyh, H.J., Cuenca, M., Field, C.M., Coelho, L.P., Cruaud, C., Engelen, S., et al. (2019). Gene Expression Changes and Community Turnover Differentially Shape the Global Ocean Metatranscriptome. Cell 179, 1068–1083 e1021.

28. Huerta-Cepas, J., Szklarczyk, D., Heller, D., Hernández-Plaza, A., Forslund, S.K., Cook, H., Mende, D.R., Letunic, I., Rattei, T., and Jensen, L.J. (2019). eggNOG 5.0: a hierarchical, functionally and phylogenetically annotated orthology resource based on 5090 organisms and 2502 viruses. Nucleic acids research 47, D309–D314.

29. Kanehisa, M., and Goto, S. (2000). KEGG: kyoto encyclopedia of genes and genomes. Nucleic acids research 28, 27–30.

30. Bahram, M., Hildebrand, F., Forslund, S.K., Anderson, J.L., Soudzilovskaia, N.A., Bodegom, P.M., Bengtsson-Palme, J., Anslan, S., Coelho, L.P., Harend, H., et al. (2018). Structure and function of the global topsoil microbiome. Nature 560, 233–237.

31. Ovchinnikov, S., Park, H., Varghese, N., Huang, P.-S., Pavlopoulos, G.A., Kim, D.E., Kamisetty, H., Kyrpides, N.C., and Baker, D. (2017). Protein structure determination using metagenome sequence data. Science 355, 294–298.

32. Laiolo, E., Alam, I., Uludag, M., Jamil, T., Agusti, S., Gojobori, T., Acinas, S.G., Gasol, J.M., and Duarte, C.M. (2024). Metagenomic probing toward an atlas of the taxonomic and metabolic foundations of the global ocean genome. Frontiers in Science 1.

33. Coelho, L.P., Alves, R., del Río, Á.R., Myers, P.N., Cantalapiedra, C.P., Giner-Lamia, J., Schmidt, T.S., Mende, D.R., Orakov, A., Letunic, I., et al. (2021). Towards the biogeography of prokaryotic genes. Nature 601, 252–256.

34. Yang, X., Garnier, F., Débat, H., Strick, T.R., and Nadal, M. (2020). Direct observation of helicase–topoisomerase coupling within reverse gyrase. Proceedings of the National Academy of Sciences 117, 10856–10864.

35. Zhang, Z., Gong, Y., Guo, L., Jiang, T., and Huang, L. (2010). Structural insights into the interaction of the crenarchaeal chromatin protein Cren7 with DNA. Molecular Microbiology 76, 749–759.

36. Boiteux, S., and Radicella, J.P. (1999). Base excision repair of 8-hydroxyguanine protects DNA from endogenous oxidative stress. Biochimie 81, 59–67.

37. Aertsen, A., De Spiegeleer, P., Vanoirbeek, K., Lavilla, M., and Michiels Chris, W. (2005). Induction of Oxidative Stress by High Hydrostatic Pressure in Escherichia coli. Applied and Environmental Microbiology 71, 2226–2231.

38. Xiao, X., Zhang, Y., and Wang, F. (2021). Hydrostatic pressure is the universal key driver of microbial evolution in the deep ocean and beyond. Environ Microbiol Rep 13, 68–72.

39. Karatzas, K.A.G., Wouters, J.A., Gahan, C.G.M., Hill, C., Abee, T., and Bennik, M.H.J. (2003). The CtsR regulator of Listeria monocytogenes contains a variant glycine repeat region that affects piezotolerance, stress resistance, motility and virulence. Molecular Microbiology 49, 1227–1238.

40. Xie, Z., Jian, H., Jin, Z., and Xiao, X. (2018). Enhancing the Adaptability of the Deep-Sea Bacterium Shewanella piezotolerans WP3 to High Pressure and Low Temperature by Experimental Evolution under H(2)O(2) Stress. Appl Environ Microbiol 84.

41. Mu, Y., Bian, C., Liu, R., Wang, Y., Shao, G., Li, J., Qiu, Y., He, T., Li, W., Ao, J., et al. (2021). Whole genome sequencing of a snailfish from the Yap Trench (∼7,000 m) clarifies the molecular mechanisms underlying adaptation to the deep sea. PLOS Genetics 17, e1009530.

42. Mougiakos, I., Mohanraju, P., Bosma, E.F., Vrouwe, V., Finger Bou, M., Naduthodi, M.I.S., Gussak, A., Brinkman, R.B.L., van Kranenburg, R., and van der Oost, J. (2017). Characterizing a thermostable Cas9 for bacterial genome editing and silencing. Nat Commun 8, 1647.

43. Schmidt, S.T., Yu, F.B., Blainey, P.C., May, A.P., and Quake, S.R. (2019). Nucleic acid cleavage with a hyperthermophilic Cas9 from an uncultured Ignavibacterium. Proceedings of the National Academy of Sciences 116, 23100–23105.

44. Ran, F.A., Cong, L., Yan, W.X., Scott, D.A., Gootenberg, J.S., Kriz, A.J., Zetsche, B., Shalem, O., Wu, X., Makarova, K.S., et al. (2015). In vivo genome editing using Staphylococcus aureus Cas9. Nature 520, 186–191.

45. Komor, A.C., Badran, A.H., and Liu, D.R. (2017). CRISPR-Based Technologies for the Manipulation of Eukaryotic Genomes. Cell 169, 559.

46. Reysenbach, A.L., St John, E., Meneghin, J., Flores, G.E., Podar, M., Dombrowski, N., Spang, A., L’Haridon, S., Humphris, S.E., de Ronde, C.E.J., et al. (2020). Complex subsurface hydrothermal fluid mixing at a submarine arc volcano supports distinct and highly diverse microbial communities. Proc Natl Acad Sci U S A 117, 32627–32638.

47. Ohta, T. (1992). The Nearly Neutral Theory of Molecular Evolution. Annual Review of Ecology and Systematics 23, 263–286.

48. Nielsen, R. (2005). Molecular signatures of natural selection. Annu Rev Genet 39, 197–218.

49. Eyre-Walker, A., and Keightley, P.D. (2007). The distribution of fitness effects of new mutations. Nat Rev Genet 8, 610–618.

50. Shapiro, B.J., and Alm, E.J. (2008). Comparing patterns of natural selection across species using selective signatures. PLoS Genet 4, e23.

51. Shapiro, B.J., and Alm, E. (2009). The slow:fast substitution ratio reveals changing patterns of natural selection in gamma-proteobacterial genomes. ISME J 3, 1180–1192.

52. Hermisson, J., and Pennings, P.S. (2005). Soft sweeps: molecular population genetics of adaptation from standing genetic variation. Genetics 169, 2335–2352.

53. Barrett, R.D., and Schluter, D. (2008). Adaptation from standing genetic variation. Trends Ecol Evol 23, 38–44.

54. Mira, A., Ochman, H., and Moran, N.A. (2001). Deletional bias and the evolution of bacterial genomes. Trends in Genetics 17, 589–596.

55. Kuo, C.-H., and Ochman, H. (2009). Deletional Bias across the Three Domains of Life. Genome Biology and Evolution 1, 145–152.

56. Bobay, L.-M., and Ochman, H. (2017). The Evolution of Bacterial Genome Architecture. Frontiers in Genetics Volume 8 - 2017.

57. Illergard, K., Ardell, D.H., and Elofsson, A. (2009). Structure is three to ten times more conserved than sequence--a study of structural response in protein cores. Proteins 77, 499–508.

58. Nomburg, J., Doherty, E.E., Price, N., Bellieny-Rabelo, D., Zhu, Y.K., and Doudna, J.A. (2024). Birth of protein folds and functions in the virome. Nature 633, 710–717.

59. Zhang, S., Zhang, T., and Fu, Y. (2023). Proteome-wide structural analysis quantifies structural conservation across distant species. Genome Res 33, 1975–1993.

60. Xu, J., and Zhang, Y. (2010). How significant is a protein structure similarity with TM-score = 0.5? Bioinformatics 26, 889–895.

61. Barrio-Hernandez, I., Yeo, J., Jänes, J., Mirdita, M., Gilchrist, C.L.M., Wein, T., Varadi, M., Velankar, S., Beltrao, P., and Steinegger, M. (2023). Clustering predicted structures at the scale of the known protein universe. Nature 622, 637–645.

62. Burley, S.K., Berman, H.M., Kleywegt, G.J., Markley, J.L., Nakamura, H., and Velankar, S. (2017). Protein Data Bank (PDB): the single global macromolecular structure archive. Protein crystallography: methods and protocols, 627–641.

63. Nogueira, T., Touchon, M., and Rocha, E.P.C. (2012). Rapid Evolution of the Sequences and Gene Repertoires of Secreted Proteins in Bacteria. PLOS ONE 7, e49403.

64. van Kempen, M., Kim, S.S., Tumescheit, C., Mirdita, M., Lee, J., Gilchrist, C.L.M., Soding, J., and Steinegger, M. (2024). Fast and accurate protein structure search with Foldseek. Nat Biotechnol 42, 243–246.

65. Lau, A.M., Bordin, N., Kandathil, S.M., Sillitoe, I., Waman, V.P., Wells, J., Orengo, C.A., and Jones, D.T. (2024). Exploring structural diversity across the protein universe with The Encyclopedia of Domains. Science 386, eadq4946.

66. Tai, C.H., Paul, R., Dukka, K.C., Shilling, J.D., and Lee, B. (2014). SymD webserver: a platform for detecting internally symmetric protein structures. Nucleic Acids Res 42, W296–300.

67. Dong, X., Peng, Y., Wang, M., Woods, L., Wu, W., Wang, Y., Xiao, X., Li, J., Jia, K., Greening, C., et al. (2023). Evolutionary ecology of microbial populations inhabiting deep sea sediments associated with cold seeps. Nature Communications 14, 1127.

68. Roda-Garcia, J.J., Haro-Moreno, J.M., and López-Pérez, M. (2023). Evolutionary pathways for deep-sea adaptation in marine planktonic Actinobacteriota. Frontiers in Microbiology 14, 1159270.

69. Craig, J.M., Laszlo, A.H., Brinkerhoff, H., Derrington, I.M., Noakes, M.T., Nova, I.C., Tickman, B.I., Doering, K., de Leeuw, N.F., and Gundlach, J.H. (2017). Revealing dynamics of helicase translocation on single-stranded DNA using high-resolution nanopore tweezers. Proceedings of the National Academy of Sciences 114, 11932–11937.

70. Garalde, D.R., Snell, E.A., Jachimowicz, D., Sipos, B., Lloyd, J.H., Bruce, M., Pantic, N., Admassu, T., James, P., and Warland, A. (2018). Highly parallel direct RNA sequencing on an array of nanopores. Nature methods 15, 201–206.

71. Dorey, A., and Howorka, S. (2024). Nanopore DNA sequencing technologies and their applications towards single-molecule proteomics. Nature chemistry 16, 314–334.

72. Fairman-Williams, M.E., Guenther, U.-P., and Jankowsky, E. (2010). SF1 and SF2 helicases: family matters. Current opinion in structural biology 20, 313–324.

73. Raney, K.D., Byrd, A.K., and Aarattuthodiyil, S. (2013). Structure and mechanisms of SF1 DNA helicases. DNA helicases and DNA motor proteins, 17–46.

74. O’Leary, N.A., Wright, M.W., Brister, J.R., Ciufo, S., Haddad, D., McVeigh, R., Rajput, B., Robbertse, B., Smith-White, B., Ako-Adjei, D., et al. (2016). Reference sequence (RefSeq) database at NCBI: current status, taxonomic expansion, and functional annotation. Nucleic Acids Res 44, D733–745.

75. Hernández-Plaza, A., Szklarczyk, D., Botas, J., Cantalapiedra, C.P., Giner-Lamia, J., Mende, D.R., Kirsch, R., Rattei, T., Letunic, I., and Jensen, L.J. (2023). eggNOG 6.0: enabling comparative genomics across 12 535 organisms. Nucleic Acids Research 51, D389–D394.

76. Byrd, A.K., and Raney, K.D. (2017). Structure and function of Pif1 helicase. Biochemical Society Transactions 45, 1159–1171.

77. Saikrishnan, K., Griffiths, S.P., Cook, N., Court, R., and Wigley, D.B. (2008). DNA binding to RecD: role of the 1B domain in SF1B helicase activity. The EMBO journal 27, 2222–2229.

78. Saikrishnan, K., Powell, B., Cook, N.J., Webb, M.R., and Wigley, D.B. (2009). Mechanistic basis of 5′-3′ translocation in SF1B helicases. Cell 137, 849–859.

79. He, X., Byrd, A.K., Yun, M.-K., Pemble, C.W., Harrison, D., Yeruva, L., Dahl, C., Kreuzer, K.N., Raney, K.D., and White, S.W. (2012). The T4 phage SF1B helicase Dda is structurally optimized to perform DNA strand separation. Structure 20, 1189–1200.

80. Saito, M., Xu, P., Faure, G., Maguire, S., Kannan, S., Altae-Tran, H., Vo, S., Desimone, A., Macrae, R.K., and Zhang, F. (2023). Fanzor is a eukaryotic programmable RNA-guided endonuclease. Nature 620, 660–668.

81. Huang, J., Lin, Q., Fei, H., He, Z., Xu, H., Li, Y., Qu, K., Han, P., Gao, Q., Li, B., et al. (2024). Discovery of deaminase functions by structure-based protein clustering. Cell 187, 4426–4428.

82. Xu, K., Feng, H., Zhang, H., He, C., Kang, H., Yuan, T., Shi, L., Zhou, C., Hua, G., Cao, Y., et al. (2025). Structure-guided discovery of highly efficient cytidine deaminases with sequence-context independence. Nat Biomed Eng 9, 93–108.

83. Zhang, C., Shine, M., Pyle, A.M., and Zhang, Y. (2022). US-align: universal structure alignments of proteins, nucleic acids, and macromolecular complexes. Nat Methods 19, 1109–1115.

84. Nji, E., Cramer, K.C., Ruffin, N.V., Fofana, F.G., Heiba, W., and Sankhe, S. (2025). Leveraging AlphaFold for innovation and sustainable health research in Africa. Nat Commun 16, 1334.

85. Yeo, J., Han, Y., Bordin, N., Lau, A.M., Kandathil, S.M., Kim, H., Karin, E.L., Mirdita, M., Jones, D.T., Orengo, C., et al. (2025). Metagenomic-scale analysis of the predicted protein structure universe. bioRxiv, 2025.2004.2023.650224.

86. Huber, C., and Wachtershauser, G. (1997). Activated acetic acid by carbon fixation on (Fe,Ni)S under primordial conditions. Science 276, 245–247.

87. Lee, S., Kim, G., Karin, E.L., Mirdita, M., Park, S., Chikhi, R., Babaian, A., Kryshtafovych, A., and Steinegger, M. (2023). Petascale Homology Search for Structure Prediction. bioRxiv.

88. Jing, X., Wu, F., Luo, X., and Xu, J. (2024). Single-sequence protein structure prediction by integrating protein language models. Proc Natl Acad Sci U S A 121, e2308788121.

89. Hayes, T., Rao, R., Akin, H., Sofroniew, N.J., Oktay, D., Lin, Z., Verkuil, R., Tran, V.Q., Deaton, J., Wiggert, M., et al. (2025). Simulating 500 million years of evolution with a language model. Science 387, 850–858.

90. Brixi, G., Durrant, M.G., Ku, J., Poli, M., Brockman, G., Chang, D., Gonzalez, G.A., King, S.H., Li, D.B., Merchant, A.T., et al. (2025). Genome modeling and design across all domains of life with Evo 2. bioRxiv.

91. Chen, S., Zhou, Y., Chen, Y., and Gu, J. (2018). fastp: an ultra-fast all-in-one FASTQ preprocessor. Bioinformatics 34, i884–i890.

92. Li, D., Luo, R., Liu, C.M., Leung, C.M., Ting, H.F., Sadakane, K., Yamashita, H., and Lam, T.W. (2016). MEGAHIT v1.0: A fast and scalable metagenome assembler driven by advanced methodologies and community practices. Methods 102, 3–11.

93. Zhu, W., Lomsadze, A., and Borodovsky, M. (2010). Ab initio gene identification in metagenomic sequences. Nucleic acids research 38, e132–e132.

94. Zhu, W., Lomsadze, A., and Borodovsky, M. (2010). Ab initio gene identification in metagenomic sequences. Nucleic Acids Res 38, e132.

95. Steinegger, M., and Soding, J. (2017). MMseqs2 enables sensitive protein sequence searching for the analysis of massive data sets. Nat Biotechnol 35, 1026–1028.

96. Uritskiy, G.V., DiRuggiero, J., and Taylor, J. (2018). MetaWRAP-a flexible pipeline for genome-resolved metagenomic data analysis. Microbiome 6, 158.

97. Parks, D.H., Imelfort, M., Skennerton, C.T., Hugenholtz, P., and Tyson, G.W. (2015). CheckM: assessing the quality of microbial genomes recovered from isolates, single cells, and metagenomes. Genome research 25, 1043–1055.

98. Olm, M.R., Brown, C.T., Brooks, B., and Banfield, J.F. (2017). dRep: a tool for fast and accurate genomic comparisons that enables improved genome recovery from metagenomes through de-replication. The ISME journal 11, 2864–2868.

99. Chaumeil, P.A., Mussig, A.J., Hugenholtz, P., and Parks, D.H. (2019). GTDB-Tk: a toolkit to classify genomes with the Genome Taxonomy Database. Bioinformatics 36, 3.

100. Parks, D.H., Chuvochina, M., Waite, D.W., Rinke, C., Skarshewski, A., Chaumeil, P.A., and Hugenholtz, P. (2018). A standardized bacterial taxonomy based on genome phylogeny substantially revises the tree of life. Nat Biotechnol 36, 996–1004.

101. Buchfink, B., Xie, C., and Huson, D.H. (2015). Fast and sensitive protein alignment using DIAMOND. Nat Methods 12, 59–60.

102. Cantalapiedra, C.P., Hernandez-Plaza, A., Letunic, I., Bork, P., and Huerta-Cepas, J. (2021). eggNOG-mapper v2: Functional Annotation, Orthology Assignments, and Domain Prediction at the Metagenomic Scale. Mol Biol Evol 38, 5825–5829.

103. Kanehisa, M., Furumichi, M., Sato, Y., Matsuura, Y., and Ishiguro-Watanabe, M. (2025). KEGG: biological systems database as a model of the real world. Nucleic Acids Res 53, D672–D677.

104. Mistry, J., Chuguransky, S., Williams, L., Qureshi, M., Salazar, G.A., Sonnhammer, E.L.L., Tosatto, S.C.E., Paladin, L., Raj, S., Richardson, L.J., et al. (2021). Pfam: The protein families database in 2021. Nucleic Acids Res 49, D412–D419.

105. Mistry, J., Finn, R.D., Eddy, S.R., Bateman, A., and Punta, M. (2013). Challenges in homology search: HMMER3 and convergent evolution of coiled-coil regions. Nucleic Acids Res 41, e121.

106. Hopf, T.A., Ingraham, J.B., Poelwijk, F.J., Scharfe, C.P., Springer, M., Sander, C., and Marks, D.S. (2017). Mutation effects predicted from sequence co-variation. Nat Biotechnol 35, 128–135.

107. Shen, W., Le, S., Li, Y., and Hu, F. (2016). SeqKit: A Cross-Platform and Ultrafast Toolkit for FASTA/Q File Manipulation. PLoS One 11, e0163962.

108. Tully, B.J., Graham, E.D., and Heidelberg, J.F. (2018). The reconstruction of 2,631 draft metagenome-assembled genomes from the global oceans. Sci Data 5, 170203.

109. Hyatt, D., LoCascio, P.F., Hauser, L.J., and Uberbacher, E.C. (2012). Gene and translation initiation site prediction in metagenomic sequences. Bioinformatics 28, 2223–2230.

110. Li, H. (2018). Minimap2: pairwise alignment for nucleotide sequences. Bioinformatics 34, 3094–3100.

111. Coelho, L.P., Alves, R., Monteiro, P., Huerta-Cepas, J., Freitas, A.T., and Bork, P. (2019). NG-meta-profiler: fast processing of metagenomes using NGLess, a domain-specific language. Microbiome 7, 1–10.

112. Fisher, R.A. (1950). Statistical methods for research workers, 11th edn (New York: Hafner).

113. Satija, R., Farrell, J.A., Gennert, D., Schier, A.F., and Regev, A. (2015). Spatial reconstruction of single-cell gene expression data. Nature biotechnology 33, 495–502.

114. Cramér, H. (1946). Mathematical methods of statistics (Princeton: Princeton university press).

115. UniProt, C. (2025). UniProt: the Universal Protein Knowledgebase in 2025. Nucleic Acids Res 53, D609–D617.

116. Katoh, K., and Standley, D.M. (2013). MAFFT multiple sequence alignment software version 7: improvements in performance and usability. Molecular biology and evolution 30, 772–780.

117. Wong, T.K., Ly-Trong, N., Ren, H., Baños, H., Roger, A.J., Susko, E., Bielow, C., De Maio, N., Goldman, N., and Hahn, M.W. (2025). IQ-TREE 3: Phylogenomic Inference Software using Complex Evolutionary Models.

118. Vasimuddin, M., Misra, S., Li, H., and Aluru, S. (2019). Efficient Architecture-Aware Acceleration of BWA-MEM for Multicore Systems. Paper presented at: 2019 IEEE International Parallel and Distributed Processing Symposium (IPDPS).

119. Danecek, P., Bonfield, J.K., Liddle, J., Marshall, J., Ohan, V., Pollard, M.O., Whitwham, A., Keane, T., McCarthy, S.A., Davies, R.M., et al. (2021). Twelve years of SAMtools and BCFtools. GigaScience 10.

120. Thorvaldsdóttir, H., Robinson, J.T., and Mesirov, J.P. (2012). Integrative Genomics Viewer (IGV): high-performance genomics data visualization and exploration. Briefings in Bioinformatics 14, 178–192.

121. Clement, K., Rees, H., Canver, M.C., Gehrke, J.M., Farouni, R., Hsu, J.Y., Cole, M.A., Liu, D.R., Joung, J.K., Bauer, D.E., et al. (2019). CRISPResso2 provides accurate and rapid genome editing sequence analysis. Nat Biotechnol 37, 224–226.

122. Smith, M.D., Wertheim, J.O., Weaver, S., Murrell, B., Scheffler, K., and Kosakovsky Pond, S.L. (2015). Less Is More: An Adaptive Branch-Site Random Effects Model for Efficient Detection of Episodic Diversifying Selection. Molecular Biology and Evolution 32, 1342–1353.

123. Edgar, R.C. (2004). MUSCLE: multiple sequence alignment with high accuracy and high throughput. Nucleic acids research 32, 1792–1797.

124. Suyama, M., Torrents, D., and Bork, P. (2006). PAL2NAL: robust conversion of protein sequence alignments into the corresponding codon alignments. Nucleic acids research 34, W609–W612.

125. Price, M.N., Dehal, P.S., and Arkin, A.P. (2010). FastTree 2--approximately maximum-likelihood trees for large alignments. PLoS One 5, e9490.

126. Kosakovsky Pond, S.L., Poon, A.F., Velazquez, R., Weaver, S., Hepler, N.L., Murrell, B., Shank, S.D., Magalis, B.R., Bouvier, D., and Nekrutenko, A. (2020). HyPhy 2.5—a customizable platform for evolutionary hypothesis testing using phylogenies. Molecular biology and evolution 37, 295–299.

127. Capella-Gutiérrez, S., Silla-Martínez, J.M., and Gabaldón, T. (2009). trimAl: a tool for automated alignment trimming in large-scale phylogenetic analyses. Bioinformatics 25, 1972–1973.

128. Langmead, B., and Salzberg, S.L. (2012). Fast gapped-read alignment with Bowtie 2. Nature methods 9, 357–359.

129. Olm, M.R., Crits-Christoph, A., Bouma-Gregson, K., Firek, B.A., Morowitz, M.J., and Banfield, J.F. (2021). inStrain profiles population microdiversity from metagenomic data and sensitively detects shared microbial strains. Nature Biotechnology 39, 727–736.

130. Benjamini, Y., and Hochberg, Y. (1995). Controlling the false discovery rate: a practical and powerful approach to multiple testing. Journal of the Royal statistical society: series B (Methodological) 57, 289–300.

131. Camacho, C., Coulouris, G., Avagyan, V., Ma, N., Papadopoulos, J., Bealer, K., and Madden, T.L. (2009). BLAST+: architecture and applications. BMC bioinformatics 10, 1–9.

132. Zhang, Y., and Skolnick, J. (2005). TM-align: a protein structure alignment algorithm based on the TM-score. Nucleic Acids Res 33, 2302–2309.

133. Abramson, J., Adler, J., Dunger, J., Evans, R., Green, T., Pritzel, A., Ronneberger, O., Willmore, L., Ballard, A.J., and Bambrick, J. (2024). Accurate structure prediction of biomolecular interactions with AlphaFold 3. Nature 630, 493–500.

134. Zhang, Y., and Skolnick, J. (2004). Scoring function for automated assessment of protein structure template quality. Proteins: Structure, Function, and Bioinformatics 57, 702–710.

135. Van Kempen, M., Kim, S.S., Tumescheit, C., Mirdita, M., Lee, J., Gilchrist, C.L., Söding, J., and Steinegger, M. (2024). Fast and accurate protein structure search with Foldseek. Nature biotechnology 42, 243–246.

136. Krogh, A., Larsson, B., von Heijne, G., and Sonnhammer, E.L. (2001). Predicting transmembrane protein topology with a hidden Markov model: application to complete genomes. J Mol Biol 305, 567–580.

137. Teufel, F., Almagro Armenteros, J.J., Johansen, A.R., Gislason, M.H., Pihl, S.I., Tsirigos, K.D., Winther, O., Brunak, S., von Heijne, G., and Nielsen, H. (2022). SignalP 6.0 predicts all five types of signal peptides using protein language models. Nat Biotechnol 40, 1023–1025.

138. Lau, A.M., Kandathil, S.M., and Jones, D.T. (2023). Merizo: a rapid and accurate protein domain segmentation method using invariant point attention. Nat Commun 14, 8445.

139. Wells, J., Hawkins-Hooker, A., Bordin, N., Sillitoe, I., Paige, B., and Orengo, C. (2024). Chainsaw: protein domain segmentation with fully convolutional neural networks. Bioinformatics 40.

140. Zhu, K., Su, H., Peng, Z., and Yang, J. (2023). A unified approach to protein domain parsing with inter-residue distance matrix. Bioinformatics 39.

141. Kandathil, S.M., Lau, A.M., Buchan, D.W.A., and Jones, D.T. (2024). Foldclass and Merizo-search: embedding-based deep learning tools for protein domain segmentation, fold recognition and comparison. bioRxiv.

142. Johnson, L.S., Eddy, S.R., and Portugaly, E. (2010). Hidden Markov model speed heuristic and iterative HMM search procedure. BMC Bioinformatics 11, 431.

143. Zhang, C., Shine, M., Pyle, A.M., and Zhang, Y. (2022). US-align: universal structure alignments of proteins, nucleic acids, and macromolecular complexes. Nature methods 19, 1109–1115.

144. Jongeneel, C., Formosa, T., and Alberts, B.M. (1984). Purification and characterization of the bacteriophage T4 dda protein. A DNA helicase that associates with the viral helix-destabilizing protein. Journal of Biological Chemistry 259, 12925–12932.

145. Winter, G., Lobley, C.M., and Prince, S.M. (2013). Decision making in xia2. Acta Crystallographica Section D: Biological Crystallography 69, 1260–1273.

146. McCoy, A.J., Grosse-Kunstleve, R.W., Adams, P.D., Winn, M.D., Storoni, L.C., and Read, R.J. (2007). Phaser crystallographic software. Journal of applied crystallography 40, 658–674.

147. Adams, P.D., Afonine, P.V., Bunkóczi, G., Chen, V.B., Davis, I.W., Echols, N., Headd, J.J., Hung, L.-W., Kapral, G.J., and Grosse-Kunstleve, R.W. (2010). PHENIX: a comprehensive Python-based system for macromolecular structure solution. Acta Crystallographica Section D: Biological Crystallography 66, 213–221.

148. Emsley, P., Lohkamp, B., Scott, W.G., and Cowtan, K. (2010). Features and development of Coot. Acta Crystallographica Section D: Biological Crystallography 66, 486–501.

149. Zhang, J.-Y., Zhang, Y., Wang, L., Guo, F., Yun, Q., Zeng, T., Yan, X., Yu, L., Cheng, L., Wu, W., et al. (2024). A single-molecule nanopore sequencing platform. bioRxiv, 2024.2008.2019.608720.

