## Supplemental Figure 1-4, Supplemental Table 2-4 for "The genetic repertoire of the deep sea: from sequence to structure and function"

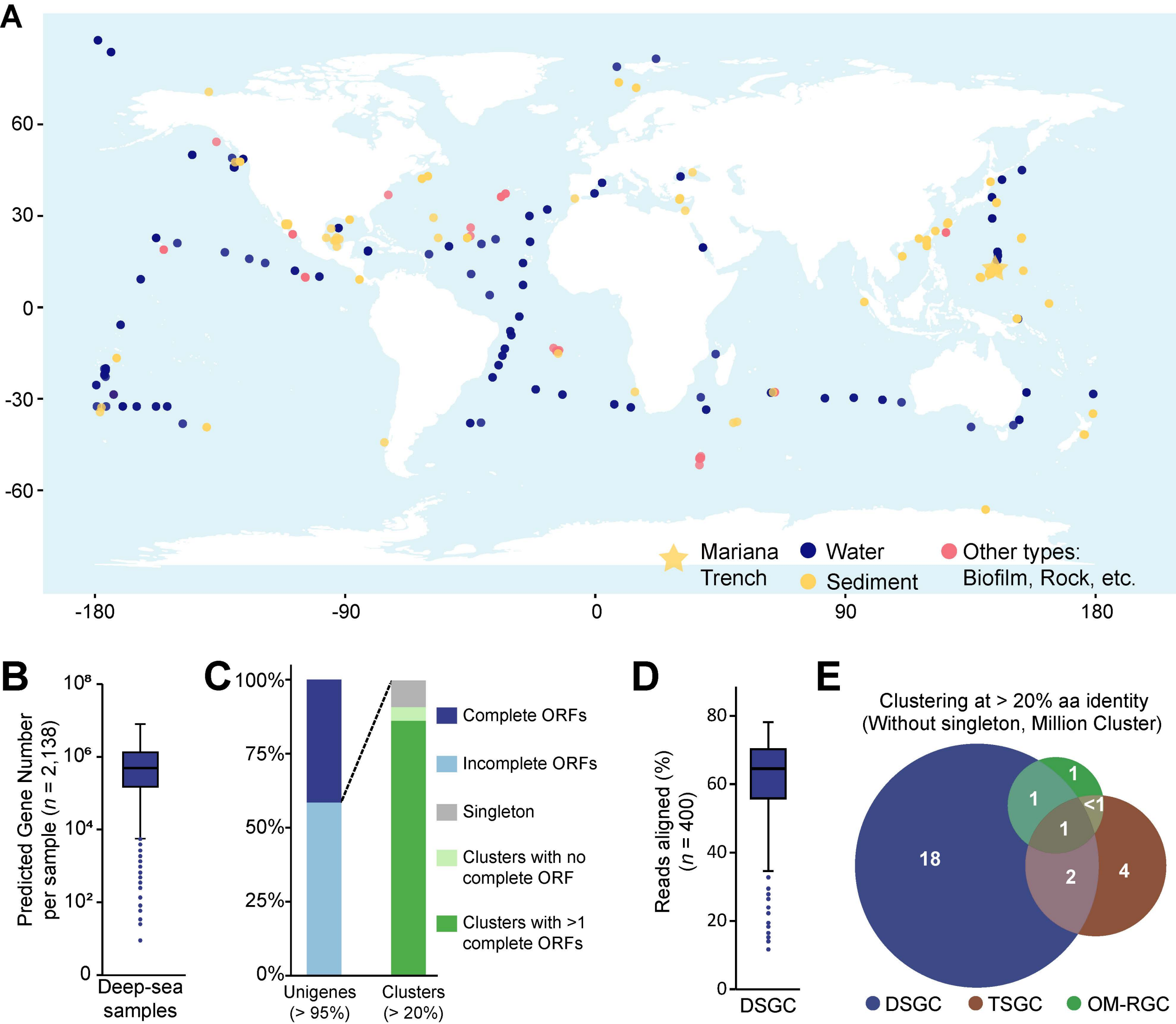
A
60
30
0
-30
-60
-180
-90
0
90
180
Mariana Trench
Water
Sediment
Other types: Biofilm, Rock, etc.
B
Predicted Gene Number per sample (n = 2,138)
10⁸
10⁶
10⁴
10²
0
Deep-sea samples
C
100%
75%
50%
25%
0%
Unigenes (> 95%)
Clusters (> 20%)
Complete ORFs
Incomplete ORFs
Singleton
Clusters with no complete ORF
Clusters with >1 complete ORFs
D
Reads aligned (%) (n = 400)
80
60
40
20
0
DSGC
E
Clustering at > 20% aa identity
(Without singleton, Million Cluster)
1
1
<1
1
18
2
4
DSGC
TSGC
OM-RGC

**Figure S1. Sample constitution and quality control of DSGC.**

**(A)** Sample types of the 2,138 deep-sea samples in this study.

**(B)** Numbers of predicted genes from the metagenomes in the 2,138 deep-sea samples.

**(C)** Clustering relationships between complete and incomplete open reading frames (ORFs) in DSGC. The amino acid identity threshold for clustering of unigenes and clusters is shown in the brackets (i.e., 95% and 20%, respectively).

**(D)** Distribution of read-mapping rates of 400 randomly selected samples out of the 2,138 deep-sea samples.

**(E)** Venn diagram of unique and shared gene clusters (at > 20% amino-acid identity with singleton cluster removed, in millions) among DSGC, OM-RGC, and TSGC.

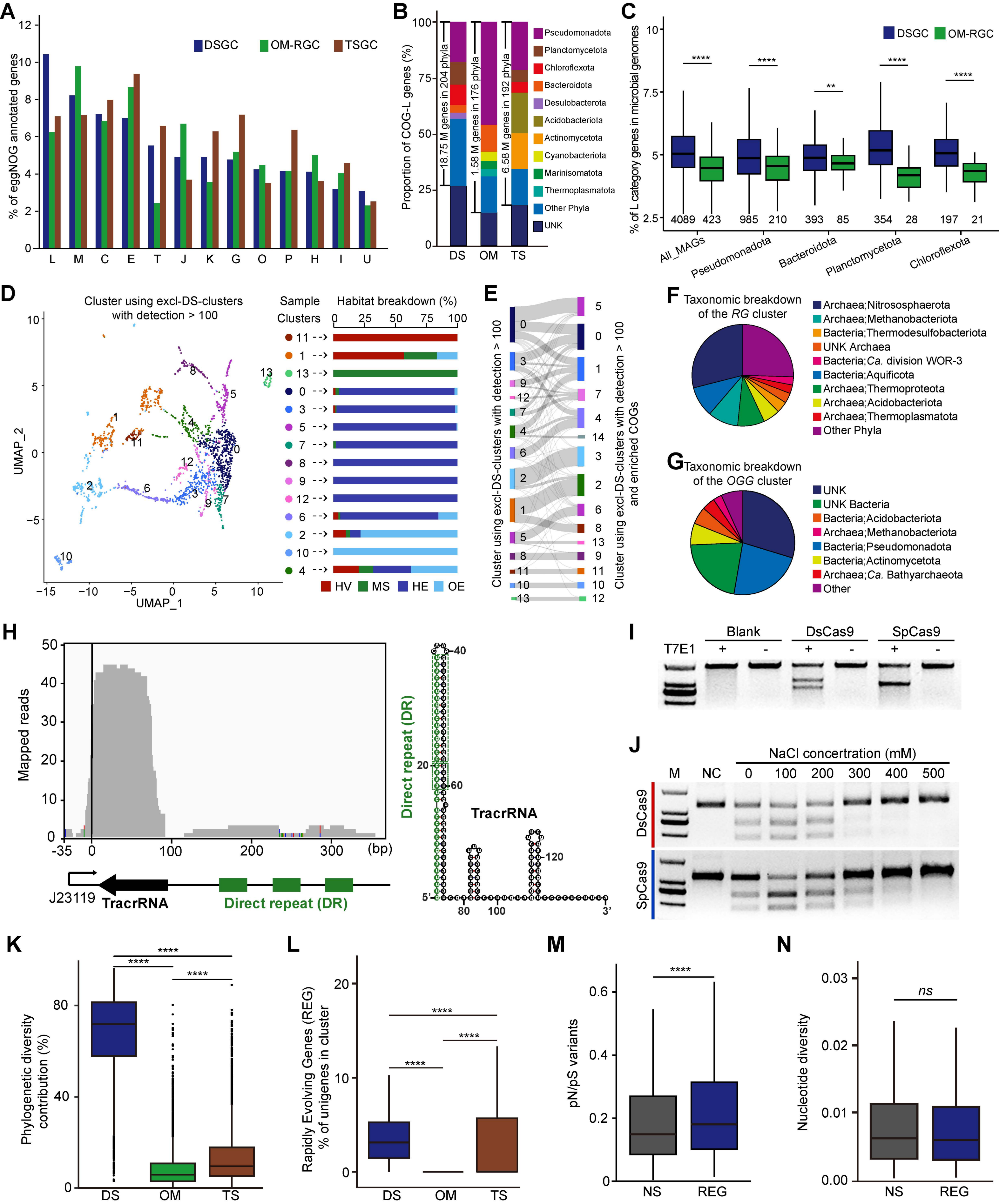
A
% of eggNOG annotated genes
DSGC
OM-RGC
TSGC
L M C E T J K G O P H I U
B
Proportion of COG-L genes (%)
18.75 M genes in 204 phyla
1.58 M genes in 176 phyla
6.58 M genes in 192 phyla
DS OM TS
Pseudomonadota
Planctomycetota
Chloroflexota
Bacteroidota
Desulobacterota
Acidobacteriota
Actinomycetota
Cyanobacteriota
Marinisomatota
Thermoplasmatota
Other Phyla
UNK
C
% of L category genes in microbial genomes
DSGC
OM-RGC
4089 423 985 210 393 85 354 28 197 21
All_MAGs
Pseudomonadota
Bacteroidota
Planctomycetota
Chloroflexota
D
Cluster using excl-DS-clusters with detection > 100
UMAP_1
UMAP_2
Sample Clusters
Habitat breakdown (%)
HV MS HE OE
E
Cluster using excl-DS-clusters with detection > 100
Cluster using excl-DS-clusters with detection > 100 and enriched COGs
F
Taxonomic breakdown of the RG cluster
Archaea;Nitrososphaerota
Archaea;Methanobacteriota
Bacteria;Thermodesulfobacteriota
UNK Archaea
Bacteria;Ca. division WOR-3
Bacteria;Aquificota
Archaea;Thermoproteota
Archaea;Acidobacteriota
Archaea;Thermoplasmatota
Other Phyla
G
Taxonomic breakdown of the OGG cluster
UNK
UNK Bacteria
Bacteria;Acidobacteriota
Archaea;Methanobacteriota
Bacteria;Pseudomonadota
Bacteria;Actinomycetota
Archaea;Ca. Bathyarchaeota
Other
H
Mapped reads
(bp)
J23119
TracrRNA
Direct repeat (DR)
5'
3'
I
T7E1
Blank
DsCas9
SpCas9
J
NaCl concertration (mM)
M NC 0 100 200 300 400 500
DsCas9
SpCas9
K
Phylogenetic diversity contribution (%)
DS OM TS
L
Rapidly Evolving Genes (REG) % of unigenes in cluster
DS OM TS
M
pN/pS variants
NS REG
N
Nucleotide diversity
ns
NS REG

**Figure S2. Diversity, distribution and evolution of COG-L genes in the deep sea.**

**(A)** Comparison of COG category constitutions among DSGC, OM-RGC, and TSGC. The categories representing < 3% genes in DSGC and the category "Function Unknown, S" are not shown. The abbreviations of categories are as follows: L, Replication, recombination and repair; M, Cell wall/membrane/envelope biogenesis; C, Energy production and conversion; E, Amino acid transport and metabolism; T, Signal transduction mechanisms; J, Translation; ribosomal structure and biogenesis; K, Transcription; G, Carbohydrate transport and metabolism; O, Posttranslational modification, protein turnover, chaperones; P, Inorganic ion transport and metabolism; H, Coenzyme transport and metabolism; I, Lipid transport and metabolism; U, Intracellular trafficking, secretion, and vesicular transport.

**(B)** Taxonomic breakdown of COG-L genes in DSGC, TSGC, and OM-RGC. Numbers of taxonomically annotatable COG-L genes and the according numbers of the annotated phyla are marked beside the columns.

**(C)** Proportion of COG-L genes in microbial genomes from the deep sea and the upper ocean. Microbial genomes were generated using the datasets of DSGC and OM-RGC, respectively. The number of genomes in each group is indicated below each box, only phyla with more than 20 high-quality MAGs from either DSGC or OM-RGC datasets were compared.

**(D)** UMAP visualization of the abundances of all prevalent excl-DS-clusters from 2,138 samples and habitat breakdown of samples in each cluster. HV: hydrothermal vents, MS: methane seeps, HE: hadal ecosystems, and OE: other deep-sea ecosystems.

**(E)** Sankey diagram of samples clustered with all prevalent excl-DS-clusters and those with enriched COG functional terms.

**(F)** Taxonomic breakdown of the hydrothermal vent-specific *RG* (COG1110) cluster. UNK: unknown taxa.

**(G)** Taxonomic breakdown of the hadal ecosystem-specific *OGG* (COG4047) cluster. UNK: unknown taxa.

**(H)** Statistics of small RNA sequencing data (left) and predicted topology structures of DsCas9's guide RNA (right).

**(I)** Human genome editing activities of DsCas9 and SpCas9 (positive control).

**(J)** *In-vitro* genome editing activities of DsCas9 and SpCas9 under different ion concentrations. M: marker, NC: negative control.

**(K)** Phylogenetic diversity contributed by DSGC, OM-RGC, and TSGC in each gene cluster.

**(L)** Frequencies of rapidly evolving genes (REGs) in each gene cluster. Selection test employed L-INS-i and IQ-Tree3 for multi-sequence alignment and tree construction, respectively. Fractions were calculated separately for genes from DSGC, OM-RGC, and TSGC, respectively. Significances of paired comparisons were assessed by one-tailed Wilcoxon test (DSGC > OM-RGC, DSGC > TSGC, TSGC > OM-RGC; ****$P < 0.0001$).

**(M)** pN/pS ratios of REGs and non-rapidly evolving genes (genes with no significant probability,

NS).

**(N)** InStrain's nucleotide diversity of REGs and NSs.

**(B, K, J)** DS:DSGC, OM: OM-RGC, TS: TSGC. **(C, K–N)** significances of paired comparisons were calculated using a two-tailed Wilcoxon test (**$P < 0.01$, ****$P < 0.0001$).

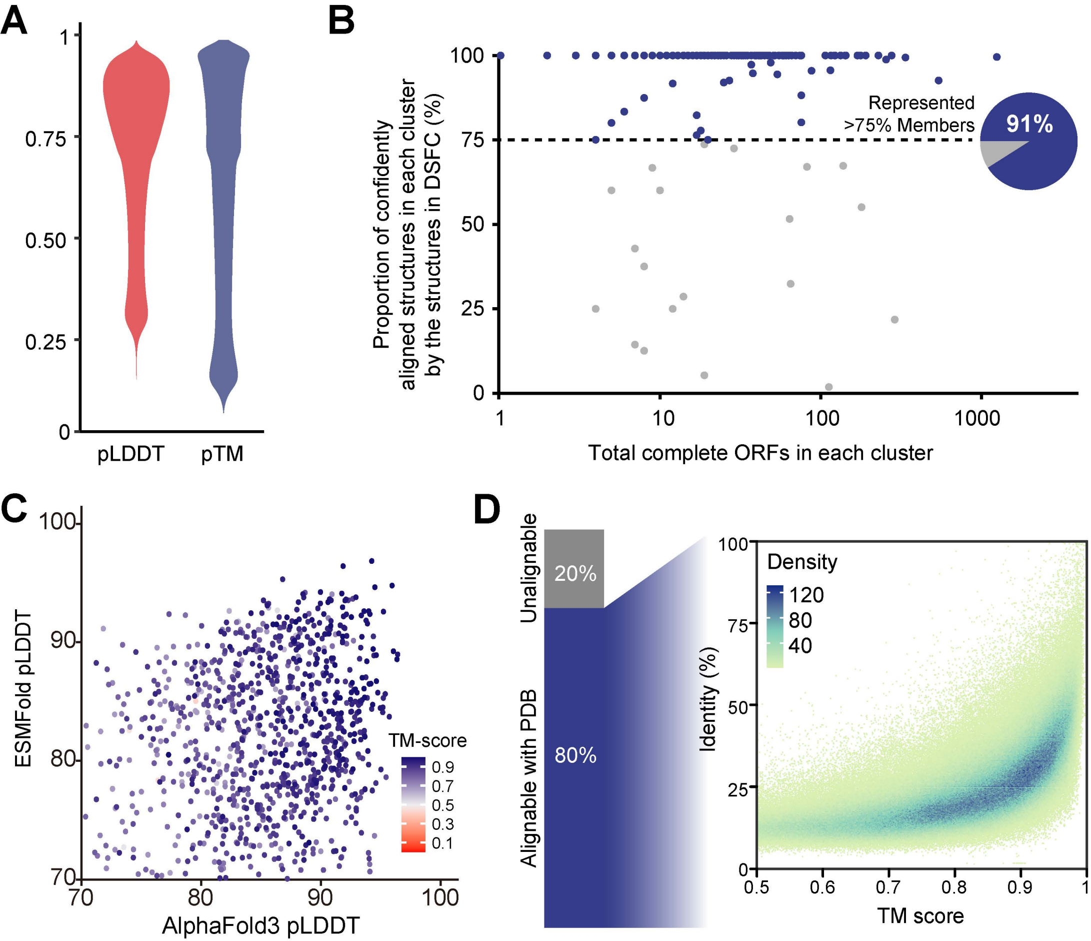

A
1
0.75
0.50
0.25
0
pLDDT
pTM
B
Proportion of confidently aligned structures in each cluster by the structures in DSFC (%)
100
75
50
25
0
Represented >75% Members
91%
1
10
100
1000
Total complete ORFs in each cluster
C
ESMFold pLDDT
100
90
80
70
TM-score
0.9
0.7
0.5
0.3
0.1
70
80
90
100
AlphaFold3 pLDDT
D
Unalignable
20%
Alignable with PDB
80%
Identity (%)
100
75
50
25
0
Density
120
80
40
0.5
0.6
0.7
0.8
0.9
1
TM score

**Figure S3. Quality control of DSFC.**

**(A)** Distribution of mean pLDDT and pTM of all structures in DSFC.

**(B)** Representativeness of structure in DSFC to other protein structures in the same sequence cluster (at > 20% amino-acid identity). 10,156 high-confidence structures from 220 clusters were predicted and compared for this analysis.

**(C)** Consistency analysis of structure predictions using ESMFold and AlphaFold3 (AF3) for deep-sea proteins ($n$ = 1,000). AF3 predicted 963 out of 1,000 high confidence (pLDDT > 70) structures, and 99.69% (960/963) of these models exhibited the same fold (TM-score > 0.5) with those predicted by ESMFold. Dots are colored by TM-scores between AF3- and ESMFold-predicted models.

**(D)** Distribution of structural and sequence similarity between proteins from DSFC and PDB.

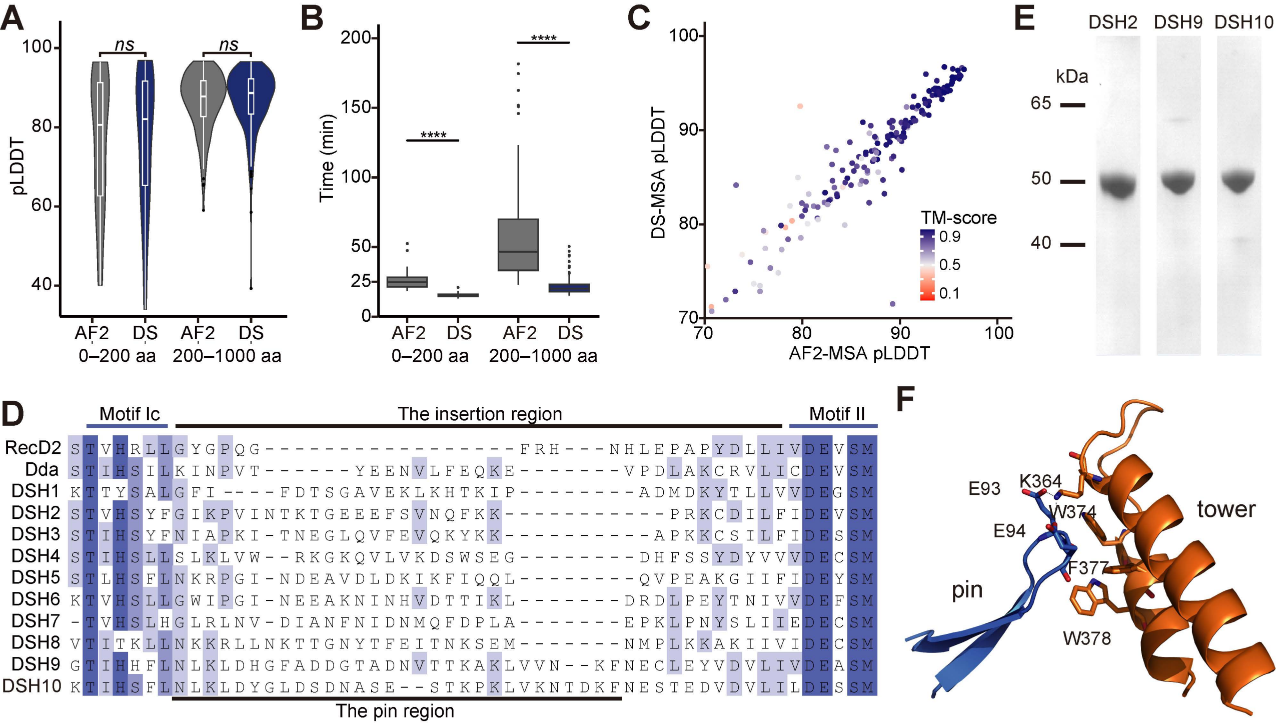

A
pLDDT
100
80
60
40
ns
ns
AF2 DS AF2 DS
0–200 aa
200–1000 aa
B
Time (min)
200
150
100
50
25
0
****
****
AF2 DS AF2 DS
0–200 aa
200–1000 aa
C
DS-MSA pLDDT
AF2-MSA pLDDT
70 80 90 100
TM-score
0.9
0.5
0.1
E
DSH2 DSH9 DSH10
kDa
65
50
40
D
Motif Ic
The insertion region
Motif II
RecD2 STVHRLLGYGPQG------------------FRH---NHLEPAPYDLLIVDEVSM
Dda STIHSILKINPVT------YEENVLFEQKE--------VPDLAKCRVLICDEVSM
DSH1 KTTYSALGFI----FDTSGAVEKLKHTKIP---------ADMDKYTLLVVDEGSM
DSH2 STVHSYFGIKPVINTKTGEEEFSVNQFKK-------------PRKCDILFIDEVSM
DSH3 STIHSYFNIAPKI-TNEGLQVFEVQKYKK------------APKKCSILFIDESSM
DSH4 STIHSLLSLKLVW--RKGKQVLVKDSWSEG--------DHFSSYDYVVVDECSM
DSH5 STLHSFLNKRPGI-NDEAVDLDKIKFIQQL---------QVPEAKGIIFIDEYSM
DSH6 KTVHSLLGWIPGI-NEEAKNINHVDTTIKL--------DRDLPEYTNIVVDEFSM
DSH7 -TVHSLHGLRLNV-DIANFNIDNMQFDPLA--------EPKLPNYSLIIEDECSM
DSH8 VTITKLLNKKRLLNKTTGNYTFEITNKSEM------NMPLKKAKIIVIDECSM
DSH9 GTIHHFLNLKLDHGFADDGTADNVTTKAKLVVN--KFNECLEYVDVLIVDEASM
DSH10 KTIHSFLNLKLDYGLDSDNASE--STKPKLVKNTDKFNESTEDVDVLILDESSM
The pin region
F
E93
K364
W374
tower
E94
F377
pin
W378

**Figure S4 Performance of DSGC-optimized AlphaFold2 and expression of helicases.**

**(A)** Prediction accuracies for 200 randomly sequences chosen from DSGC using multiple-sequence-alignments (MSAs) with the AlphaFold2 (AF2) default metagenomic datasets and DSGC respectively. Statistics for shorter sequences (0–200 aa, $n = 40$) and longer sequences (200–1000 aa, $n = 160$) were computed separately. The DSGC-optimized AF2 showed comparable accuracies with the default AlphaFold2 in predicting deep-sea structures.

**(B)** Ratios of computing time of the structure prediction using the AF2 default metagenomic datasets (1.8 TB) and DSGC (95 GB). The use of DSGC-MSAs resulted in a sharp reduction in computing time for predictions.

**(C)** Consistency analysis of structure predictions using AF2 with DS-MSA and AF2-MSA for deep-sea proteins ($n = 200$). Dots are colored by TM-scores between models generated using DS-MSA and AF2-MSA.

**(D)** MSA of the pin-like 1B domain of DSH1–10, in comparison to those of RecD2 and Dda.

**(E)** The SDS-PAGE image of purified DSH2, DSH9, and DSH10.

**(F)** Key interactions between the pin and the tower in Dda (PDB: 3UPU).

**(A, B)** significances of paired comparisons were calculated using two-tailed Wilcoxon test (ns: no significant, ****$P < 0.0001$).

**Table S1 Metadata of metagenome datasets used in this study[1-80]**

**Table S2 Clustering statistics from five independent replicates of random subsampling (10 million genes per catalog)**

| | Shared by all catalogs | Shared by DS[a] and TS[b] | Shared by DS and OM[c] | Shared by TS and OM | DS exclusive | TS exclusive | OM exclusive | DS exclusive without singletons | TS exclusive without singletons | OM exclusive without singletons |
|---|---|---|---|---|---|---|---|---|---|---|
| **Repeat 1** | 184,677 | 176,711 | 179,483 | 50,494 | 3,408,552 | 2,577,954 | 1,431,624 | 465,353 | 402,836 | 319,525 |
| **Repeat 2** | 184,556 | 176,931 | 179,778 | 50,826 | 3,406,256 | 2,579,356 | 1,432,089 | 465,495 | 402,629 | 319,650 |
| **Repeat 3** | 184,474 | 176,649 | 179,250 | 50,872 | 3,407,154 | 2,581,017 | 1,431,407 | 466,522 | 402,851 | 319,309 |
| **Repeat 4** | 184,981 | 176,458 | 179,066 | 50,924 | 3,407,840 | 2,578,251 | 1,430,437 | 465,187 | 403,817 | 319,613 |
| **Repeat 5** | 184,740 | 176,807 | 179,337 | 50,681 | 3,405,358 | 2,579,271 | 1,430,780 | 466,413 | 403,081 | 319,055 |
| **Average** | 184,686 | 176,711 | 179,383 | 50,759 | 3,407,032 | 2,579,170 | 1,431,267 | 465,794 | 403,043 | 319,430 |

[a]**DS: DSGC**

[b]**TS: TSGC**

[c]**OM: OM-RGC**

**Table S3 Numbers of non-redundant genes assigned to the overlapping taxonomic groups in each catalog**

| Taxa | | OM-RGC | TSGC | DSGC |
|---|---|---|---|---|
| **Phylum-level** | | | | |
| **d_Bacteria;p_Pseudomonadota**[a] | | 17,523,184 | 34,953,158 | 64,444,507 |
| **d_Bacteria;p_Chloroflexota** | | 794,658 | 6,747,097 | 36,186,910 |
| **d_Bacteria;p_Planctomycetota** | | 828,942 | 5,839,647 | 35,886,553 |
| **d_Bacteria;p_Bacteroidota** | | 4,914,206 | 5,267,143 | 17,287,337 |
| **d_Bacteria;p_Desulfobacterota** | | 51,518 | 559,735 | 10,812,818 |
| **d_Bacteria;p_Patescibacteriota** | | 287,201 | 502,080 | 9,595,012 |
| **d_Archaea;p_Thermoproteota** | | 405,920 | 887,525 | 9,205,497 |
| **d_Bacteria;p_Gemmatimonadota** | | 88,233 | 2,635,795 | 7,006,275 |
| **d_Bacteria;p_Bacteroidota_A** | | 130,122 | 188,292 | 3,120,526 |
| **d_Bacteria;p_Myxococcota** | | 228,095 | 2,280,342 | 3,106,005 |
| **d_Bacteria;p_Bacillota** | | 113,603 | 851,672 | 2,799,186 |
| d_Bacteria;p_Acidobacteriota | | 168,146 | 29,387,919 | 8,705,759 |
| d_Bacteria;p_Actinomycetota | | 783,470 | 23,599,735 | 5,579,446 |
| d_Bacteria;p_Verrucomicrobiota | | 1,134,796 | 5,775,926 | 4,240,088 |
| d_Bacteria;p_Cyanobacteriota | | 1,563,120 | 417,252 | 974,880 |
| | sum | 29,015,214 | 119,893,318 | 218,950,799 |
| **Order-level** | | | | |
| **d_Bacteria;p_Chloroflexota; c_Dehalococcoidia;o_UBA3495** | | 145,215 | 16,870 | 9,669,500 |
| **d_Bacteria;p_Planctomycetota; c_Phycisphaerae;o_Phycisphaerales** | | 182,432 | 81,861 | 6,355,783 |
| **d_Archaea;p_Thermoproteota; c_Nitrososphaeria;o_Nitrososphaerales** | | 392,621 | 708,830 | 6,241,270 |
| **d_Bacteria;p_Bacteroidota; c_Bacteroidia;o_Flavobacteriales** | | 3,905,426 | 363,176 | 5,869,910 |
| **d_Bacteria;p_Bacteroidota; c_Bacteroidia;o_Bacteroidales** | | 92,307 | 266,647 | 5,741,371 |
| **d_Bacteria;p_Planctomycetota; c_Planctomycetia;o_Pirellulales** | | 327,653 | 767,779 | 5,677,495 |
| **d_Bacteria;p_Planctomycetota; c_Phycisphaerae;o_Sedimentisphaerales** | | 6,364 | 21,208 | 5,167,406 |
| **d_Bacteria;p_Pseudomonadota; c_Gammaproteobacteria;o_Pseudomonadales** | | 2,086,547 | 968,501 | 5,044,763 |
| **d_Bacteria;p_Myxococcota_A; c_UBA9160;o_UBA9160** | | 29,457 | 84,258 | 3,845,057 |

| | | | |
|---|---|---|---|
| **d_Bacteria;p_Pseudomonadota; c_Alphaproteobacteria;o_Rhodobacterales** | 1,082,686 | 76,822 | 3,764,380 |
| **d_Bacteria;p_Desulfobacterota; c_Desulfobacteria;o_Desulfobacterales** | 22,450 | 34,101 | 3,646,467 |
| **d_Bacteria;p_Gemmatimonadota; c_Gemmatimonadetes;o_Longimicrobiales** | 71,665 | 102,066 | 3,403,143 |
| **d_Bacteria;p_Pseudomonadota; c_Alphaproteobacteria;o_Rhodospirillales** | 370,678 | 91,698 | 3,273,303 |
| **d_Bacteria;p_Gemmatimonadota; c_Gemmatimonadetes;o_Gemmatimonadales** | 8,836 | 2,494,359 | 2,573,170 |
| **d_Bacteria;p_Chloroflexota; c_Anaerolineae;o_Anaerolineales** | 44,235 | 606,728 | 2,546,156 |
| **d_Bacteria;p_Planctomycetota; c_Planctomycetia;o_Planctomycetales** | 118,426 | 384,278 | 2,451,025 |
| **d_Bacteria;p_Pseudomonadota; c_Gammaproteobacteria;o_Enterobacterales** | 613,645 | 175,831 | 2,166,567 |
| **d_Bacteria;p_Pseudomonadota; c_Gammaproteobacteria;o_Xanthomonadales** | 15,885 | 809,208 | 1,871,011 |
| **d_Bacteria;p_Bacteroidota; c_Bacteroidia;o_Cytophagales** | 226,548 | 467,129 | 1,853,738 |
| **d_Bacteria;p_Patescibacteriota; c_Minisyncoccia;o_UBA9973** | 83,721 | 88,153 | 1,607,161 |
| **d_Bacteria;p_Desulfobacterota; c_Desulfobulbia;o_Desulfobulbales** | 5,203 | 36,994 | 1,590,675 |
| **d_Bacteria;p_Bacteroidota_A; c_Ignavibacteria;o_Ignavibacteriales** | 10,715 | 51,519 | 1,525,390 |
| **d_Bacteria;p_Pseudomonadota; c_Gammaproteobacteria;o_Methylococcales** | 20,506 | 30,340 | 1,288,068 |
| **d_Bacteria;p_Planctomycetota; c_UBA1135;o_UBA1135** | 32,395 | 43,223 | 1,054,155 |
| **d_Bacteria;p_Pseudomonadota; c_Gammaproteobacteria;o_Chromatiales** | 7,409 | 18,892 | 940,438 |
| **d_Bacteria;p_Nitrospirota; c_Nitrospiria;o_Nitrospirales** | 10,906 | 259,375 | 818,207 |
| **d_Bacteria;p_Myxococcota; c_Polyangia;o_Polyangiales** | 16,786 | 798,692 | 816,079 |
| **d_Bacteria;p_Verrucomicrobiota; c_Verrucomicrobiia;o_Opitutales** | 491,007 | 314,332 | 746,646 |
| **d_Bacteria;p_Verrucomicrobiota; c_Verrucomicrobiia;o_Verrucomicrobiales** | 234,897 | 191,635 | 651,994 |
| **d_Bacteria;p_SAR324; c_SAR324;o_SAR324** | 188,941 | 47,790 | 506,925 |
| **d_Bacteria;p_Chlamydiota; c_Chlamydiia;o_Chlamydiales** | 21,283 | 149,747 | 404,861 |
| **d_Bacteria;p_Cyanobacteriota; c_Cyanobacteriia;o_Cyanobacteriales** | 14,911 | 176,880 | 380,715 |
| **d_Bacteria;p_Babelota; c_Babeliae;o_Babelales** | 7,842 | 39,543 | 352,828 |

| | | | |
|---|---|---|---|
| **d_Bacteria;p_Bacillota;<br>c_Clostridia;o_Lachnospirales** | 12,087 | 23,900 | 351,056 |
| **d_Bacteria;p_Bdellovibrionota;<br>c_Bacteriovoracia;o_Bacteriovoracales** | 135,860 | 20,435 | 337,484 |
| **d_Bacteria;p_Myxococcota;<br>c_UBA796;o_UBA796** | 64,767 | 35,843 | 309,842 |
| **d_Bacteria;p_Pseudomonadota;<br>c_Gammaproteobacteria;o_Rariloculales** | 26,426 | 64,511 | 293,066 |
| **d_Bacteria;p_Pseudomonadota;<br>c_Alphaproteobacteria;o_Rickettsiales** | 36,058 | 26,378 | 188,292 |
| **d_Bacteria;p_Bdellovibrionota;<br>c_Bdellovibrionia;o_Bdellovibrionales** | 32,571 | 50,397 | 144,674 |
| **d_Bacteria;p_Pseudomonadota;<br>c_Gammaproteobacteria;o_Nevskiales** | 33,034 | 37,051 | 96,948 |
| **d_Bacteria;p_Chloroflexota;<br>c_Dehalococcoidia;o_Tepidiformales** | 11,022 | 24,411 | 86,753 |
| **d_Bacteria;p_Bacteroidota;<br>c_Bacteroidia;o_2-12-FULL-35-15** | 20,037 | 27,351 | 63,152 |
| **d_Bacteria;p_Myxococcota;<br>c_UBA727;o_UBA727** | 11,664 | 22,315 | 59,719 |
| **d_Bacteria;p_Pseudomonadota;<br>c_Gammaproteobacteria;o_Legionellales** | 19,064 | 32,755 | 55,563 |
| **d_Bacteria;p_Bacillota;<br>c_Clostridia;o_Clostridiales** | 12,199 | 50,358 | 54,457 |
| d_Bacteria;p_Pseudomonadota;<br>c_Alphaproteobacteria;o_Rhizobiales | 197,782 | 13,122,475 | 3,556,709 |
| d_Bacteria;p_Pseudomonadota;<br>c_Gammaproteobacteria;o_Burkholderiales | 377,601 | 5,394,821 | 1,971,299 |
| d_Bacteria;p_Bacteroidota;<br>c_Bacteroidia;o_Chitinophagales | 287,648 | 2,736,764 | 1,547,691 |
| d_Bacteria;p_Pseudomonadota;<br>c_Alphaproteobacteria;o_Sphingomonadales | 278,831 | 1,610,845 | 1,534,986 |
| d_Bacteria;p_Actinomycetota;<br>c_Acidimicrobiia;o_Acidimicrobiales | 392,288 | 1,693,716 | 1,118,556 |
| d_Bacteria;p_Pseudomonadota;<br>c_Alphaproteobacteria;o_Caulobacterales | 402,326 | 1,097,338 | 1,040,041 |
| d_Bacteria;p_Acidobacteriota;<br>c_Vicinamibacteria;o_Vicinamibacterales | 111,551 | 3,859,641 | 1,034,205 |
| d_Bacteria;p_Verrucomicrobiota;<br>c_Verrucomicrobiia;o_Limisphaerales | 325,865 | 1,300,495 | 827,386 |
| d_Bacteria;p_Acidobacteriota;<br>c_Terriglobia;o_Bryobacterales | 6,785 | 4,247,609 | 552,894 |
| d_Bacteria;p_Actinomycetota;<br>c_Actinomycetes;o_Mycobacteriales | 16,110 | 4,468,065 | 445,633 |
| d_Bacteria;p_Actinomycetota;<br>c_Actinomycetes;o_Actinomycetales | 65,985 | 1,239,496 | 332,790 |
| d_Bacteria;p_Myxococcota;<br>c_Polyangia;o_Haliangiales | 10,487 | 367,193 | 260,823 |

| | | | |
|---|---|---|---|
| d_Bacteria;p_Patescibacteriota; c_Saccharimonadia;o_Saccharimonadales | 23,821 | 198,415 | 177,600 |
| d_Bacteria;p_Actinomycetota; c_Actinomycetes;o_Propionibacteriales | 11,100 | 1,324,701 | 158,198 |
| d_Bacteria;p_Bacteroidota; c_Bacteroidia;o_Sphingobacteriales | 8,887 | 791,415 | 110,753 |
| d_Bacteria;p_Verrucomicrobiota; c_Verrucomicrobiia;o_Chthoniobacterales | 7,570 | 3,654,236 | 97,183 |
| d_Bacteria;p_Bacteroidota; c_Bacteroidia;o_NS11-12g | 77,765 | 17,539 | 72,233 |
| sum | 13,906,789 | 58,308,934 | 110,725,643 |

[a]Bolded taxa indicates DSGC's gene number is the largest among all three catalogs.

**Table S4 Data collection and refinement statistics**

| | DSH9 |
|---|---|
| **Data collection** | |
| Space group | $P2_12_12_1$ |
| Cell dimensions | |
| *a*, *b*, *c* (Å) | 56.81 126.84 150.71 |
| α, β, γ (°) | 90.00 90.00 90.00 |
| Resolution (Å)[a] | 49.08-2.06 (2.17-2.06) |
| Reflections (total/unique) | 673482/63166 |
| $R_{merge}$[b] | 0.056 (0.485) |
| CC1/2[c] | 99.9 (87.6) |
| I/σ(I) | 23.1 (3.0) |
| Completeness (%) | 92.5 (86.6) |
| Redundancy | 10.7(5.9) |
| **Refinement** | |
| $R_{work/}$ $R_{free}$ | 0.200/0.245 |
| R.m.s deviations | |
| Bond lengths (Å) | 0.002 |
| Bond angles (º) | 0.419 |
| Ramachandran (%)[d] plot(%) | 97.31/2.69/0.00 |
| Residue range | A/1-111,119-454; B/4-454 |
| H2O | 655 |
| Other ligand | 2 Tartrates |
| Metal ion | 2 MG |

[a]Values for highest resolution shells are given in parentheses.
[b]Rmerge = $\sum hkl \sum_i |I_i(hkl) - \langle I(hkl)\rangle| / \sum hkl \sum_i |I_i(hkl)$ where $I_i$ (*hkl*) and $\langle I(hkl)\rangle$ are the i and mean measurement of intensity of reflection *hkl.*
[c]CC1/2 = Pearson's correlation coefficient between average intensities of random half data sets for each unique reflection.
[d]Residues in favored, accepted, and outlier regions of the Ramachandran plot as reported by MOLPROBITY

### References


1. Anantharaman, K., Duhaime, M.B., Breier, J.A., Wendt, K.A., Toner, B.M., and Dick, G.J. (2014). Sulfur oxidation genes in diverse deep-sea viruses. Science *344*, 757-760.
2. Bäckström, D., Yutin, N., Jørgensen, S.L., Dharamshi, J., Homa, F., Zaremba-Niedwiedzka, K., Spang, A., Wolf, Y.I., Koonin, E.V., and Ettema, T.J. (2019). Virus genomes from deep sea sediments expand the ocean megavirome and support independent origins of viral gigantism. MBio *10*, 10.1128/mbio. 02497-02418.
3. Beckmann, S., Farag, I.F., Zhao, R., Christman, G.D., Prouty, N.G., and Biddle, J.F. (2021). Expanding the repertoire of electron acceptors for the anaerobic oxidation of methane in carbonates in the Atlantic and Pacific Ocean. The ISME Journal *15*, 2523-2536.
4. Bergauer, K., Fernandez-Guerra, A., Garcia, J.A., Sprenger, R.R., Stepanauskas, R., Pachiadaki, M.G., Jensen, O.N., and Herndl, G.J. (2018). Organic matter processing by microbial communities throughout the Atlantic water column as revealed by metaproteomics. Proceedings of the National Academy of Sciences *115*, E400-E408.
5. Biller, S.J., Berube, P.M., Dooley, K., Williams, M., Satinsky, B.M., Hackl, T., Hogle, S.L., Coe, A., Bergauer, K., Bouman, H.A.*, et al.* (2018). Marine microbial metagenomes sampled across space and time. Scientific Data *5*, 180176.
6. Boeuf, D., Edwards, B.R., Eppley, J.M., Hu, S.K., Poff, K.E., Romano, A.E., Caron, D.A., Karl, D.M., and DeLong, E.F. (2019). Biological composition and microbial dynamics of sinking particulate organic matter at abyssal depths in the oligotrophic open ocean. Proceedings of the National Academy of Sciences *116*, 11824-11832.
7. Bowman, K.L., Collins, R.E., Agather, A.M., Lamborg, C.H., Hammerschmidt, C.R., Kaul, D., Dupont, C.L., Christensen, G.A., and Elias, D.A. (2020). Distribution of mercury-cycling genes in the Arctic and equatorial Pacific Oceans and their relationship to mercury speciation. Limnology and Oceanography *65*, S310-S320.
8. Coe, A., Ghizzoni, J., LeGault, K., Biller, S., Roggensack, S.E., and Chisholm, S.W. (2016). Survival of Prochlorococcus in extended darkness. Limnology and Oceanography *61*, 1375-1388.
9. Coutinho, F.H., von Meijenfeldt, F.A.B., Walter, J.M., Haro-Moreno, J.M., Lopéz-Pérez, M., van Verk, M.C., Thompson, C.C., Cosenza, C.A.N., Appolinario, L., Paranhos, R.*, et al.* (2021). Ecogenomics and metabolic potential of the South Atlantic Ocean microbiome. Science of The Total Environment *765*, 142758.
10. Defforey, D. (2016). Phosphorus cycling in the deep sedimentary subseafloor environment (University of California, Santa Cruz).
11. Dombrowski, N., Seitz, K.W., Teske, A.P., and Baker, B.J. (2017). Genomic insights into potential interdependencies in microbial hydrocarbon and nutrient cycling in hydrothermal sediments. Microbiome *5*.
12. Dombrowski, N., Teske, A.P., and Baker, B.J. (2018). Expansive microbial metabolic versatility and biodiversity in dynamic Guaymas Basin hydrothermal sediments. Nature communications *9*, 4999.
13. Dong, X., Rattray, J.E., Campbell, D.C., Webb, J., Chakraborty, A., Adebayo, O., Matthews, S., Li, C., Fowler, M., and Morrison, N.M. (2020). Thermogenic hydrocarbon biodegradation by diverse depth-stratified microbial populations at a Scotian Basin cold seep. Nature communications *11*, 5825.
14. Fortunato, C.S., Larson, B., Butterfield, D.A., and Huber, J.A. (2018). Spatially distinct, temporally stable microbial populations mediate biogeochemical cycling at and below the seafloor in hydrothermal vent fluids. Environmental Microbiology *20*, 769-784.
15. Fullerton, H., Hager, K.W., McAllister, S.M., and Moyer, C.L. (2017). Hidden diversity revealed by genome-resolved metagenomics of iron-oxidizing microbial mats from Lō'ihi Seamount, Hawai'i. The ISME Journal *11*, 1900-1914.
16. He, T., Li, H., and Zhang, X. (2017). Deep-Sea Hydrothermal Vent Viruses Compensate for Microbial Metabolism in Virus-Host Interactions. mBio *8*, 10.1128/mbio.00893-00817.
17. He, Y., Li, M., Perumal, V., Feng, X., Fang, J., Xie, J., Sievert, S.M., and Wang, F. (2016). Genomic and enzymatic evidence for acetogenesis among multiple lineages of the archaeal phylum Bathyarchaeota widespread in marine sediments. Nature Microbiology *1*, 16035.
18. Huang, W.-C., Liu, Y., Zhang, X., Zhang, C.-J., Zou, D., Zheng, S., Xu, W., Luo, Z., Liu, F., and Li, M. (2021). Comparative genomic analysis reveals metabolic flexibility of Woesearchaeota. Nature Communications *12*, 5281.
19. Jarett, J., Dunfield, P., Peura, S., Wielen, P.v.d., Hedlund, B., Elshahed, M., Kormas, K., Birkeland, N.-K., Zhang, C., Rengefors, K.*, et al.* (2014). Microbial Dark Matter phase II: stepping deeper into unknown territory (Lawrence Berkeley National Lab.(LBNL), Berkeley, CA (United States)).
20. Jungbluth, S.P., Bowers, R.M., Lin, H.T., Cowen, J.P., and Rappe, M.S. (2016). Novel microbial assemblages inhabiting crustal fluids within mid-ocean ridge flank subsurface basalt. ISME J *10*, 2033-

2047.
21. Kato, S., Hirai, M., Ohkuma, M., and Suzuki, K. (2019). Microbial metabolisms in an abyssal ferromanganese crust from the Takuyo-Daigo Seamount as revealed by metagenomics. PLoS One *14*, e0224888.
22. Kato, S., Nakano, S., Kouduka, M., Hirai, M., Suzuki, K., Itoh, T., Ohkuma, M., and Suzuki, Y. (2019). Metabolic Potential of As-yet-uncultured Archaeal Lineages of <i>Candidatus</i> Hydrothermarchaeota Thriving in Deep-sea Metal Sulfide Deposits. Microbes and Environments *34*, 293-303.
23. Kato, S., Shibuya, T., Takaki, Y., Hirai, M., Nunoura, T., and Suzuki, K. (2018). Genome-enabled metabolic reconstruction of dominant chemosynthetic colonizers in deep-sea massive sulfide deposits. Environmental microbiology *20*, 862-877.
24. Koppers, A.A., Yamazaki, T., Geldmacher, J., Anderson, L., Beier, C., Buchs, D.M., Chen, L., Cohen, B., Deschamps, F., and Dorais, M. (2011). Louisville Seamount Trail: implications for geodynamic mantle flow models and the geochemical evolution of primary hotspots.
25. Laso-Pérez, R., Hahn, C., van Vliet Daan, M., Tegetmeyer Halina, E., Schubotz, F., Smit Nadine, T., Pape, T., Sahling, H., Bohrmann, G., Boetius, A.*, et al.* (2019). Anaerobic Degradation of Non-Methane Alkanes by "Candidatus Methanoliparia" in Hydrocarbon Seeps of the Gulf of Mexico. mBio *10*, 10.1128/mbio.01814-01819.
26. Laso-Pérez, R., Wegener, G., Knittel, K., Widdel, F., Harding, K.J., Krukenberg, V., Meier, D.V., Richter, M., Tegetmeyer, H.E., and Riedel, D. (2016). Thermophilic archaea activate butane via alkyl-coenzyme M formation. Nature *539*, 396-401.
27. Laso-Pérez, R., Wu, F., Crémière, A., Speth, D.R., Magyar, J.S., Zhao, K., Krupovic, M., and Orphan, V.J. (2023). Evolutionary diversification of methanotrophic ANME-1 archaea and their expansive virome. Nature Microbiology *8*, 231-245.
28. Lecoeuvre, A., Ménez, B., Cannat, M., Chavagnac, V., and Gérard, E. (2021). Microbial ecology of the newly discovered serpentinite-hosted Old City hydrothermal field (southwest Indian ridge). The ISME Journal *15*, 818-832.
29. Li, J., Li, L., Bai, S., Ta, K., Xu, H., Chen, S., Pan, J., Li, M., Du, M., and Peng, X. (2019). New insight into the biogeochemical cycling of methane, S and Fe above the Sulfate-Methane Transition Zone in methane hydrate-bearing sediments: A case study in the Dongsha area, South China Sea. Deep Sea Research Part I: Oceanographic Research Papers *145*, 97-108.
30. Li, L., Bai, S., Li, J., Wang, S., Tang, L., Dasgupta, S., Tang, Y., and Peng, X. (2020). Volcanic ash inputs enhance the deep-sea seabed metal-biogeochemical cycle: A case study in the Yap Trench, western Pacific Ocean. Marine Geology *430*, 106340.
31. Li, M., Baker, B.J., Anantharaman, K., Jain, S., Breier, J.A., and Dick, G.J. (2015). Genomic and transcriptomic evidence for scavenging of diverse organic compounds by widespread deep-sea archaea. Nature communications *6*, 8933.
32. Li, Y., Jing, H., Xia, X., Cheung, S., Suzuki, K., and Liu, H. (2018). Metagenomic Insights Into the Microbial Community and Nutrient Cycling in the Western Subarctic Pacific Ocean. Frontiers in Microbiology *9*.
33. Liang, R., Davidova, I.A., Teske, A., and Suflita, J.M. (2023). Evidence for the anaerobic biodegradation of higher molecular weight hydrocarbons in the Guaymas Basin. International Biodeterioration & Biodegradation *181*, 105621.
34. Liu, J., Zheng, Y., Lin, H., Wang, X., Li, M., Liu, Y., Yu, M., Zhao, M., Pedentchouk, N., Lea-Smith, D.J.*, et al.* (2019). Proliferation of hydrocarbon-degrading microbes at the bottom of the Mariana Trench. Microbiome *7*, 47.
35. Lopez-Perez, M., Haro-Moreno, J.M., Gonzalez-Serrano, R., Parras-Molto, M., and Rodriguez-Valera, F. (2017). Genome diversity of marine phages recovered from Mediterranean metagenomes: size matters. PLoS genetics *13*, e1007018.
36. McAllister Sean, M., Polson Shawn, W., Butterfield David, A., Glazer Brian, T., Sylvan Jason, B., and Chan Clara, S. (2020). Validating the Cyc2 Neutrophilic Iron Oxidation Pathway Using Meta-omics of Zetaproteobacteria Iron Mats at Marine Hydrothermal Vents. mSystems *5*, 10.1128/msystems.00553-00519.
37. Meier, D.V., Pjevac, P., Bach, W., Hourdez, S., Girguis, P.R., Vidoudez, C., Amann, R., and Meyerdierks, A. (2017). Niche partitioning of diverse sulfur-oxidizing bacteria at hydrothermal vents. The ISME Journal *11*, 1545-1558.
38. Meier, D.V., Pjevac, P., Bach, W., Markert, S., Schweder, T., Jamieson, J., Petersen, S., Amann, R., and Meyerdierks, A. (2019). Microbial metal-sulfide oxidation in inactive hydrothermal vent chimneys suggested by metagenomic and metaproteomic analyses. Environmental Microbiology *21*, 682-701.
39. Mende, D.R., Bryant, J.A., Aylward, F.O., Eppley, J.M., Nielsen, T., Karl, D.M., and DeLong, E.F. (2017). Environmental drivers of a microbial genomic transition zone in the ocean's interior. Nature Microbiology

*2*, 1367-1373.
40. Meng, C., Li, S., Fan, Q., Chen, R., Hu, Y., Xiao, X., and Jian, H. (2020). The thermo-regulated genetic switch of deep-sea filamentous phage SW1 and its distribution in the Pacific Ocean. FEMS Microbiology Letters *367*.
41. Meyer, J.L., Jaekel, U., Tully, B.J., Glazer, B.T., Wheat, C.G., Lin, H.-T., Hsieh, C.-C., Cowen, J.P., Hulme, S.M., Girguis, P.R., *et al.* (2016). A distinct and active bacterial community in cold oxygenated fluids circulating beneath the western flank of the Mid-Atlantic ridge. Scientific Reports *6*, 22541.
42. Michoud, G., Ngugi, D.K., Barozzi, A., Merlino, G., Calleja, M.L., Delgado-Huertas, A., Morán, X.A.G., and Daffonchio, D. (2021). Fine-scale metabolic discontinuity in a stratified prokaryote microbiome of a Red Sea deep halocline. The ISME Journal *15*, 2351-2365.
43. Niemann, H., Lösekann, T., De Beer, D., Elvert, M., Nadalig, T., Knittel, K., Amann, R., Sauter, E.J., Schlüter, M., and Klages, M. (2006). Novel microbial communities of the Haakon Mosby mud volcano and their role as a methane sink. Nature *443*, 854-858.
44. Polzin, J., Arevalo, P., Nussbaumer, T., Polz, M.F., and Bright, M. (2019). Polyclonal symbiont populations in hydrothermal vent tubeworms and the environment. Proceedings of the Royal Society B: Biological Sciences *286*, 20181281.
45. Raggi, L., García-Guevara, F., Godoy-Lozano, E.E., Martínez-Santana, A., Escobar-Zepeda, A., Gutierrez-Rios, R.M., Loza, A., Merino, E., Sanchez-Flores, A., Licea-Navarro, A., *et al.* (2020). Metagenomic Profiling and Microbial Metabolic Potential of Perdido Fold Belt (NW) and Campeche Knolls (SE) in the Gulf of Mexico. Frontiers in Microbiology *11*.
46. Rajala, P., Cheng, D.-Q., Rice, S.A., and Lauro, F.M. (2022). Sulfate-dependant microbially induced corrosion of mild steel in the deep sea: a 10-year microbiome study. Microbiome *10*, 1-14.
47. Reveillaud, J., Reddington, E., McDermott, J., Algar, C., Meyer, J.L., Sylva, S., Seewald, J., German, C.R., and Huber, J.A. (2016). Subseafloor microbial communities in hydrogen-rich vent fluids from hydrothermal systems along the Mid-Cayman Rise. Environmental Microbiology *18*, 1970-1987.
48. Reysenbach, A.-L., St. John, E., Meneghin, J., Flores, G.E., Podar, M., Dombrowski, N., Spang, A., L'Haridon, S., Humphris, S.E., and de Ronde, C.E. (2020). Complex subsurface hydrothermal fluid mixing at a submarine arc volcano supports distinct and highly diverse microbial communities. Proceedings of the National Academy of Sciences *117*, 32627-32638.
49. Roux, S., Hawley, A.K., Torres Beltran, M., Scofield, M., Schwientek, P., Stepanauskas, R., Woyke, T., Hallam, S.J., and Sullivan, M.B. (2014). Ecology and evolution of viruses infecting uncultivated SUP05 bacteria as revealed by single-cell- and meta-genomics. eLife *3*, e03125.
50. Saw, J.H.W., Nunoura, T., Hirai, M., Takaki, Y., Parsons, R., Michelsen, M., Longnecker, K., Kujawinski, E.B., Stepanauskas, R., Landry, Z., *et al.* (2020). Pangenomics Analysis Reveals Diversification of Enzyme Families and Niche Specialization in Globally Abundant SAR202 Bacteria. mBio *11*, 10.1128/mbio.02975-02919.
51. Shin, J.H., Eom, H., Song, W.J., and Rho, M. (2018). Integrative metagenomic and biochemical studies on rifamycin ADP-ribosyltransferases discovered in the sediment microbiome. Scientific Reports *8*, 12143.
52. Shinichi Sunagawa, L.P.C., Samuel Chaffron, Eric Karsenti, Jeroen Raes, Silvia G. Acinas, Peer Bork (2015). Structure and function of the global ocean microbiome. Science *348*, 9.
53. Skennerton, C.T., Ward, L.M., Michel, A., Metcalfe, K., Valiente, C., Mullin, S., Chan, K.Y., Gradinaru, V., and Orphan, V.J. (2015). Genomic reconstruction of an uncultured hydrothermal vent gammaproteobacterial methanotroph (family Methylothermaceae) indicates multiple adaptations to oxygen limitation. Frontiers in microbiology *6*, 1425.
54. Smith, A.R., Kieft, B., Mueller, R., Fisk, M.R., Mason, O.U., Popa, R., and Colwell, F.S. (2019). Carbon fixation and energy metabolisms of a subseafloor olivine biofilm. The ISME Journal *13*, 1737-1749.
55. Song, D., Zhang, Y., Liu, J., Zhong, H., Zheng, Y., Zhou, S., Yu, M., Todd, J.D., and Zhang, X.H. (2020). Metagenomic Insights Into the Cycling of Dimethylsulfoniopropionate and Related Molecules in the Eastern China Marginal Seas. Front Microbiol *11*, 157.
56. Spang, A., Saw, J.H., Jørgensen, S.L., Zaremba-Niedzwiedzka, K., Martijn, J., Lind, A.E., Van Eijk, R., Schleper, C., Guy, L., and Ettema, T.J. (2015). Complex archaea that bridge the gap between prokaryotes and eukaryotes. Nature *521*, 173-179.
57. St. John, E., Flores, G.E., Meneghin, J., and Reysenbach, A.L. (2019). Deep-sea hydrothermal vent metagenome-assembled genomes provide insight into the phylum Nanoarchaeota. Environmental microbiology reports *11*, 262-270.
58. Tanikawa, W., Tadai, O., Morono, Y., Hinrichs, K.-U., and Inagaki, F. (2018). Geophysical constraints on microbial biomass in subseafloor sediments and coal seams down to 2.5 km off Shimokita Peninsula, Japan. Progress in Earth and Planetary Science *5*, 1-17.
59. Trembath-Reichert, E., Butterfield, D.A., and Huber, J.A. (2019). Active subseafloor microbial communities from Mariana back-arc venting fluids share metabolic strategies across different thermal niches and taxa.

The ISME Journal *13*, 2264-2279.
60. Tsementzi, D., Wu, J., Deutsch, S., Nath, S., Rodriguez-R, L.M., Burns, A.S., Ranjan, P., Sarode, N., Malmstrom, R.R., and Padilla, C.C. (2016). SAR11 bacteria linked to ocean anoxia and nitrogen loss. Nature *536*, 179-183.
61. Tully, B.J., and Heidelberg, J.F. (2016). Potential mechanisms for microbial energy acquisition in oxic deep-sea sediments. Applied and Environmental Microbiology *82*, 4232-4243.
62. Tully, B.J., Wheat, C.G., Glazer, B.T., and Huber, J.A. (2018). A dynamic microbial community with high functional redundancy inhabits the cold, oxic subseafloor aquifer. ISME J *12*, 1-16.
63. Villanueva, L., von Meijenfeldt, F.B., Westbye, A.B., Yadav, S., Hopmans, E.C., Dutilh, B.E., and Damsté, J.S.S. (2021). Bridging the membrane lipid divide: bacteria of the FCB group superphylum have the potential to synthesize archaeal ether lipids. The ISME Journal *15*, 168-182.
64. Vuillemin, A., Wankel, S.D., Coskun, Ö.K., Magritsch, T., Vargas, S., Estes, E.R., Spivack, A.J., Smith, D.C., Pockalny, R., Murray, R.W.*, et al.* Archaea dominate oxic subseafloor communities over multimillion-year time scales. Science Advances *5*, eaaw4108.
65. Wang, F.P., Zhang, Y., Chen, Y., He, Y., Qi, J., Hinrichs, K.U., Zhang, X.X., Xiao, X., and Boon, N. (2014). Methanotrophic archaea possessing diverging methane-oxidizing and electron-transporting pathways. ISME J *8*, 1069-1078.
66. Wang, H.-l., and Sun, L. (2017). Comparative metagenomics reveals insights into the deep-sea adaptation mechanism of the microorganisms in Iheya hydrothermal fields. World Journal of Microbiology and Biotechnology *33*, 86.
67. Wang, H.-l., Zhang, J., Sun, Q.-l., Lian, C., and Sun, L. (2017). A comparative study revealed first insights into the diversity and metabolisms of the microbial communities in the sediments of Pacmanus and Desmos hydrothermal fields. PLoS one *12*, e0181048.
68. Yamamoto, M., Takaki, Y., Kashima, H., Tsuda, M., Tanizaki, A., Nakamura, R., and Takai, K. (2023). In situ electrosynthetic bacterial growth using electricity generated by a deep-sea hydrothermal vent. The ISME Journal *17*, 12-20.
69. Yanagawa, K., Nunoura, T., McAllister, S.M., Hirai, M., Breuker, A., Brandt, L., House, C.H., Moyer, C.L., Birrien, J.-L., and Aoike, K. (2013). The first microbiological contamination assessment by deep-sea drilling and coring by the D/V Chikyu at the Iheya North hydrothermal field in the Mid-Okinawa Trough (IODP Expedition 331). Frontiers in microbiology *4*, 327.
70. Yu, H., Susanti, D., McGlynn, S.E., Skennerton, C.T., Chourey, K., Iyer, R., Scheller, S., Tavormina, P.L., Hettich, R.L., Mukhopadhyay, B.*, et al.* (2018). Comparative Genomics and Proteomic Analysis of Assimilatory Sulfate Reduction Pathways in Anaerobic Methanotrophic Archaea. Front Microbiol *9*, 2917.
71. Zehnle, H., Laso-Pérez, R., Lipp, J., Riedel, D., Benito Merino, D., Teske, A., and Wegener, G. (2023). Candidatus Alkanophaga archaea from Guaymas Basin hydrothermal vent sediment oxidize petroleum alkanes. Nature Microbiology *8*, 1199-1212.
72. Zhang, H., Wang, M., Wang, H., Chen, H., Cao, L., Zhong, Z., Lian, C., Zhou, L., and Li, C. (2022). Metagenome sequencing and 768 microbial genomes from cold seep in South China Sea. Scientific Data *9*, 480.
73. Zhang, W., Sun, J., Cao, H., Tian, R., Cai, L., Ding, W., and Qian, P.-Y. (2016). Post-translational modifications are enriched within protein functional groups important to bacterial adaptation within a deep-sea hydrothermal vent environment. Microbiome *4*, 49.
74. Zhang, X., Feng, X., and Wang, F. (2016). Diversity and Metabolic Potentials of Subsurface Crustal Microorganisms from the Western Flank of the Mid-Atlantic Ridge. Frontiers in Microbiology *7*.
75. Zhang, X., Xu, W., Liu, Y., Cai, M., Luo, Z., and Li, M. (2018). Metagenomics Reveals Microbial Diversity and Metabolic Potentials of Seawater and Surface Sediment From a Hadal Biosphere at the Yap Trench. Frontiers in Microbiology *9*.
76. Zhao, J., Jing, H., Wang, Z., Wang, L., Jian, H., Zhang, R., Xiao, X., Chen, F., Jiao, N., and Zhang, Y. (2022). Novel viral communities potentially assisting in carbon, nitrogen, and sulfur metabolism in the upper slope sediments of Mariana Trench. MSystems *7*, e01358-01321.
77. Zhao, R., Hannisdal, B., Mogollon, J.M., and Jørgensen, S.L. (2019). Nitrifier abundance and diversity peak at deep redox transition zones. Scientific Reports *9*, 8633.
78. Zhou, Y.-L., Mara, P., Cui, G.-J., Edgcomb, V.P., and Wang, Y. (2022). Microbiomes in the Challenger Deep slope and bottom-axis sediments. Nature Communications *13*, 1515.
79. Zhou, Z., Tran, P.Q., Kieft, K., and Anantharaman, K. (2020). Genome diversification in globally distributed novel marine Proteobacteria is linked to environmental adaptation. The ISME Journal *14*, 2060-2077.
80. Zorz, J., Li, C., Chakraborty, A., Gittins, D.A., Surcon, T., Morrison, N., Bennett, R., MacDonald, A., and Hubert, C.R.J. (2023). SituSeq: an offline protocol for rapid and remote Nanopore 16S rRNA amplicon sequence analysis. ISME Communications *3*, 33.
